# Sex chromosome-aware GWAS reveals XY-linked agronomic architecture and breeding targets in cannabis

**DOI:** 10.64898/2026.06.18.733269

**Authors:** Mehdi Babaei, Davoud Torkamaneh

## Abstract

*Cannabis sativa* is a dioecious crop whose male plants carry a Y chromosome entirely excluded from every genome-wide association study (GWAS) published to date, despite its direct relevance to agronomic trait variation. Here, we present the first sex chromosome-aware GWAS of cannabis, integrating both ChrX and ChrY across 145 landrace accessions and 27 agronomic traits. By combining SNP-based GWAS with sex-stratified *k*-mer association analysis, we identified 16 genome-wide significant markers: eight on ChrX controlling traits including stem biomass (*CsWSL1*; phenotypic variance explained (PVE) 56.4%) and relative growth rate (CsX8; PVE 73.6%), and eight on ChrY controlling male morphology (*CsMSML1*; PVE 17.18-76.1%) and male-specific flowering (*CsMSFL1*; PVE 84.08–87.1%). *K*-mer GWAS yielded 1,838 annotated signal-trait associations representing 1,082 unique genomic windows across the pseudoautosomal, hemizygous X, and male-specific Y regions. The key loci identified here provide a genomic toolkit for marker-assisted paternal selection, flowering-time synchronization, and biomass improvement across dioecious cannabis breeding systems.

**HIGHLIGHT:** First sex chromosome-aware GWAS of cannabis integrating ChrX and ChrY identifies 16 agronomic loci including *CsMSFL1*, a near-deterministic regulator of male flowering with 87% phenotypic variance explained.

## 1. INTRODUCTION

*Cannabis sativa* L. is among humanity’s oldest cultivated plants, with origins tracing back approximately 3,000–8,000 years to East/Central Asia, where it was among the earliest domesticated crops cultivated for textile, food, and medicinal purposes (Small, 2017; Ren *et al*., 2021; Babaei *et al*., 2022, 2025). Despite its deep cultural and historical significance, cannabis has lagged behind many major crop species in genomic research. This delay stems in part from decades of prohibition that constrained fundamental scientific investigation, as well as from the species’ inherently complex biology. Its repeat-rich genome, extensive phenotypic plasticity, remarkable chemotypic diversity, and dioecious reproductive —where male and female plants are genetically distinct—have collectively posed substantial challenges for genomic analysis and breeding efforts (Lapierre *et al*., 2023; Lynch *et al*., 2025). Overall, these made male genomics both commercially consequential and systematically understudied.

The past decade has seen rapid progress in cannabis genomics, including multiple reference genome assemblies and a growing body of genome-wide association studies (GWAS) linking SNP markers to phenological, morphological, and phytochemical traits (de Ronne *et al*., 2024; de Ronne and Torkamaneh, 2025; Babaei and Torkamaneh, 2026). Without exception, these studies have been conducted on autosomes. The Y chromosome, present in every male cannabis plant and constituting nearly a third of the male haploid genome, has remained outside every GWAS panel published to date, not because it is genetically trivial, but because the reference assemblies and analytical frameworks available until recently made its inclusion technically intractable.

*Cannabis* belongs to the Cannabaceae family alongside its closest relative *Humulus* (hops), and the two genera share a remarkably conserved sex chromosome system. Both possess heteromorphic XY sex chromosomes, with *H. lupulus* sharing the same XX/XY system as *Cannabis*, while *H. japonicus* carries an XX/XY_1_Y_2_ configuration (Prentout *et al*., 2021; Razumova *et al*., 2023). Genomic in situ hybridization has revealed extensive male-specific regions (MSY) with accumulated repetitive DNA on the Y chromosomes of both genera, and 112 XY-linked genes are shared between the two species (Prentout *et al*., 2021; Babaei *et al*., 2026*a*). This conserved system represents one of the oldest documented plant sex chromosome pairs, estimated to predate the divergence of the two genera at approximately 28 to 48 Mya (Prentout *et al*., 2021; Carey *et al*., 2026), making it an exceptional model for studying sex chromosome evolution and the agronomic consequences of long-term recombination suppression.

Sexual plasticity—the ability of genetically male or female plants to alter their sexual phenotype and produce flowers of the opposite sex in response to genetic, hormonal, or environmental cues—in cannabis further underscores the evolutionary and functional significance of sex chromosomes. Several genes associated with ethylene biosynthesis, perception, and signaling, pathways known to play central roles in sex determination and floral differentiation, are located within the non-recombining region of the ChrX and are notably absent from ChrY (Monthony *et al*., 2026). This unequal genomic distribution suggests that sex chromosome differentiation has been accompanied by the accumulation of regulatory elements influencing reproductive development. Together, these findings indicate that ChrX functions not only as a carrier of sex-determining factors but also as a genomic hub for genes governing reproductive flexibility and developmental responses.

The exclusion of sex chromosomes from GWAS is not a problem unique to cannabis. A landmark re-survey of the NHGRI-EBI GWAS Catalog found that only 25% of studies reported X chromosome results and a mere 3% reported Y chromosome results, a phenomenon described as an expanding exclusionary problem (Sun *et al*., 2023; Cerezo *et al*., 2025). In dioecious plants and animals alike, the consequence is a systematic underrepresentation of male-specific genetic variation in our understanding of trait architecture (Muyle *et al*., 2021; Charlesworth and Harkess, 2024). In asparagus, male plants outyield females in commercial production, yet the Y-linked basis of sex determination was characterized only recently (Harkess *et al*., 2017; Motoki *et al*., 2022). In kiwifruit and date palm, Y-linked loci have been exploited for marker-assisted seedling sex identification, enabling breeders to select male pollinators years before first flowering (Torres *et al*., 2018; Akagi *et al*., 2019). In *H. lupulus*, X-linked sex determination has been identified as a direct target for hop breeding programs (Carey *et al*., 2026). In livestock, MSY-based haplotype networks have resolved paternal lineages and male-driven population dynamics with a resolution impossible from autosomal data alone (Felkel *et al*., 2019). The pattern is consistent across kingdoms: the male sex chromosome encodes variation of direct agronomic, evolutionary, and conservation relevance, yet it is the last to receive systematic genomic attention.

In cannabis, this blind spot is particularly consequential. Male plants are the sole pollen donors in any breeding cross and contribute 50% of seed genetics in fiber and grain programs, yet they are routinely destroyed before flowering in cannabinoid production systems (Lipson Feder *et al*., 2021). Their X-linked and Y-linked genetic architecture therefore shapes trait variation across every cannabis production system, whether or not male plants themselves are retained. In fiber hemp and seed hemp programs, male plant traits — stem morphology, branching architecture, growth rate, and flowering phenology — directly affect crossing efficiency and yield; optimizing these traits requires genomic tools for paternal line selection that have not existed until now. In cannabinoid production, the male parents used for foundation seed lots determine the sex-chromosome genetic constitution of the commercial female crop, yet breeders have been unable to genotype or select on Y-linked or X-linked variants contributing to offspring performance. The absence of such tools has also precluded marker-based approaches to male–female flowering synchronization, a critical bottleneck in open-pollinated breeding programs. The recent availability of haplotype-resolved reference assemblies for both sex chromosomes, including the Otto II assembly with phased ChrX and ChrY sequences and a characterized pseudoautosomal region (PAR) and non-pseudoautosomal region (nonPAR) boundary (Carey *et al*., 2026), makes it possible for the first time to conduct a sex chromosome-inclusive GWAS with the same rigour applied to autosomes. Building on our previous GWAS of agronomic traits across the autosomal genome cannabis landrace accessions (Babaei and Torkamaneh, 2026), we extend the analysis to both sex chromosomes by integrating SNP-based GWAS with sex-stratified *k*-mer association analysis. *K*-mer GWAS complements SNP approaches by operating reference-independently, capturing structural variants absent from SNP arrays, and has demonstrated superior resolution near causal loci in crops (Torres *et al*., 2018; Voichek and Weigel, 2020; Karikari *et al*., 2023; Lemay *et al*., 2023; Jaegle *et al*., 2025).

This study aims to characterize the genetic architecture of agronomic traits encoded on ChrX and ChrY across a diverse cannabis landrace panel, resolve the population-genetic and evolutionary structure of both sex chromosomes, and identify candidate loci with direct relevance to male plant biology and dioecious breeding programs.

## 2. MATERIALS AND METHODS

### 2.1 Plant materials

A panel of 145 *Cannabis sativa* L. landrace accessions (72 males, 73 females) representing 25 native populations from Iran was used in this study (Supplementary Table S1). Seed samples were collected from local markets and native farmers across five Köppen–Geiger climate zones (BSh: arid, steppe, hot; BSk: arid, steppe, cold; BWh: arid, desert, hot; BWk: arid, desert, cold; Cfa: temperate, without dry season, sizzling summer), spanning geographic latitudes 25°–40°N and longitudes 45°–65°E. Full details of seed sourcing, cultivation, randomized complete block design (RCBD; three blocks), greenhouse conditions are described in Babaei et al. (2024) and Babaei and Torkamaneh (2026). Seed collection and cultivation were conducted with ethics approval and license from the anti-narcotics police in Razavi Khorasan. Activities at Université Laval complied fully with Health Canada regulations under license LIC-QX0ZJC7SIP-2021.

### 2.2 Phenotypic data

Phenotypic data were derived from two prior publications on this accession panel. Phenological and morphological traits were assessed as described in Babaei et al. (2024); chemotype classification was taken directly from Babaei and Torkamaneh (2026). For the present study, a total of 27 agronomic traits were analyzed (Supplementary Table S2). Six phenological traits were recorded in days after sowing: 1) GV Point (GVP), 2) Start Flower Formation Time in Individuals (SFFI); 3) Start Flower Formation Time in 50% of the Population (SFFP); 4) Start 10% Flowering Time in Individuals (SF10I); 5) Start 10% Flowering Time in 50% of the Population (SF10P); 6) Flowering Time 50% in Individuals (FT50I). Five node and branching architecture traits comprised: 1) Number of Nodes to the Main Inflorescence (NTMI); 2) Number of Nodes to the First Lateral Shoot (NTFIS); 3) Number of Nodes on the Main Stem at Harvest (NNH); 4) Number of Nodes on the Main Inflorescence (NMI); 5) Number of Lateral Shoots (NLS). Eight growth and structural dimension traits were: 1) Length of Shortest Lateral Shoot (LSLS); 2) Length of Main Inflorescence (LMI); 3) Length of Longest Lateral Shoot (LLLS); 4) Length of Internode in the Middle Third of the Main Stem at Harvest (LIMTH); 5) Height at Harvest (HH); 6) Height to GV Point (HGV); 7) Relative Growth Rate (RGR); 8) Stem Diameter at Harvest (SDH). Eight biomass yield traits were: 1) Dry Weight of Flowers (DWF); 2) Fresh Weight of Flowers (FWF); 3) Fresh Weight of Leaves (FWL); 4) Dry Weight of Leaves (DWL); 5) Fresh Weight of Stems (FWS); 6) Dry Weight of Stems (DWS); 7) Total Fresh Weight (TFW); 8) Total Dry Weight (TDW).

Chemotype was assigned to one of four classes based on THC:CBD and CBD:(THC+CBN) dual-ratio thresholds (Babaei and Torkamaneh, 2026): THC-dominant; balanced; moderate CBD; CBD-dominant. Flowering group comprised three photoperiod-response classes (Babaei *et al*., 2026*b*): auto-flowering (day-length-independent flowering), early-flowering, and late-flowering. Plant architectural type was categorized into four morphological classes: PT1 (lateral-attached), PT2 (uniform lateral), PT3 (pyramidal), and PT4 (menorah) (Babaei *et al*., 2024). These three classifications were used exclusively as metadata for phylogenetic tree annotation and were not included as GWAS phenotypes.

### 2.3 Genotyping and SNP calling

#### 2.3.1 Genotyping and variant calling

Leaf tissue was sampled from young, healthy leaves of all individuals. Tissue was homogenized with metallic beads in a RETSCH MM 400 mixer mill (Fisher Scientific, MA, USA), and DNA was extracted using the Qiagen DNeasy® Plant Mini Kit following the manufacturer’s protocol. DNA quantity was assessed with a Qubit fluorometer (dsDNA HS assay kit; Thermo Fisher Scientific, MA, USA) and quality was verified by agarose gel electrophoresis on a random subset. All samples were normalized to 10 ng/µl before library preparation.

High-density genotyping-by-sequencing (HD-GBS) libraries were constructed using *Bfa*I as the restriction enzyme following Torkamaneh et al., (2021), at the Institut de biologie intégrative et des systèmes (IBIS), Université Laval, QC, Canada. Paired-end sequencing was performed on an Illumina NovaSeq 6000 at the Genome Québec Service and Expertise Center (CESGQ), Montréal, QC, Canada, generating approximately 400 million reads in total (∼2.7 million reads per sample).

Raw reads were processed with FastGBS v2.0 (Torkamaneh *et al*., 2020) for adapter trimming and quality filtering: adapter sequences were removed and reads shorter than 50 bp were discarded. Processed reads were aligned to the haplotype-resolved Otto II assembly of *C. sativa* (Carey et al., 2026; GenBank accession [TBD]), which provides fully phased sequences for all nine autosomes and both sex chromosomes (ChrX, 87.4 Mb; ChrY, 114 Mb), using BWA v0.7.17 (Li and Durbin, 2009). Reads with mapping quality below 30 were discarded prior to variant calling. Variant calling was performed with the Platypus pipeline (Rimmer *et al*., 2014) requiring a minimum supporting read depth of six (MINREADS = 6), yielding an initial dataset of approximately 4.5 million raw variants used as input for all downstream filtering steps.

#### 2.3.2 PAR/nonPAR boundary determination

The boundary between the pseudoautosomal region (PAR) and the non-recombining compartments (hemizygous X region (HXR), on ChrX; male-specific region (MSY), on ChrY) was established using two independent approaches. Prior to analysis, the raw variant set was minimally filtered to retain biallelic SNPs passing the FILTER field only (VCFtools v0.1.16 (Danecek *et al*., 2011); *--remove-filtered-all; --min-alleles 2*; --*max-alleles 2*), without additional quality or frequency thresholds, to maximize variant coverage for boundary determination.

First, windowed log₂(female/male) read-depth ratios were computed with DifCover (Smith *et al*., 2018) using BAM files from one representative female (MB137) and one male (MB24) accession. Coverage was restricted to 10-250× per sample, ratios were computed in windows of 2,000 valid positions (minimum segment length 500 bp), and segmentation was performed with DNAcopy (log_2_ adjustment AC = 1.25). Segment-level analysis identified the first HXR-characteristic segment on ChrX at 30.04 Mb (log_2_= +0.45) and the first MSY-characteristic segment on ChrY at 31.7 Mb (log_2_ = −1.48), with intervening segments showing transitional values near the boundary. Median log_2_ ratios per compartment were: PAR (ChrX) = −0.21 and PAR (ChrY) = +0.12, consistent with diploid coverage in both sexes; HXR = +0.42, consistent with hemizygosity in males; and MSY = −1.55, confirming male-only coverage. Extreme negative log₂ values at three short ChrY segments (range: −5.5 to −8.3) corresponded to repetitive centromeric regions with near-zero mappability and were excluded from boundary inference (Supplementary Table S3; Supplementary Fig. S1).

Second, per-individual SNP density was quantified in non-overlapping 10 kb windows using BCFtools v1.19 and VCFtools v0.1.16 (--*SNPdensity 10000*). Male–female density differences were assessed by 1,000 permutations (*p* ≤ 0.001). The proportion of significantly sex-differentiated windows was 7.3% in ChrX PAR and 7.0% in ChrY PAR, consistent with background levels reflecting free recombination. This proportion increased to 15.1% in the HXR, consistent with female diploidy versus male hemizygosity, and reached 53.5% in the MSY, confirming male-specific variant enrichment. A slight male-biased SNP density in the ChrX PAR is attributable to cross-mapping of ChrY PAR reads to the ChrX PAR reference (Supplementary Fig. S2)

Both approaches converged on a PAR/nonPAR boundary of 30 Mb on both ChrX and ChrY, consistent with the boundary of 29.3 to 35.7 Mb determined for the Otto II assembly (Carey *et al*., 2026). This boundary was applied to all downstream variant filtering and association analyses.

#### 2.3.3 Sex chromosome-aware variant filtering

Raw variants were processed with a custom sex chromosome-aware filtering and imputation pipeline XYRefiner (github.com/Mehdibabaeii/XYRefiner) across five chromosomal compartments following recommendations for sex chromosome GWAS quality control (König *et al*., 2014). After genome-wide basic filters (FILTER = PASS; biallelic; QUAL ≥ 10), region-specific quality control was applied: Hardy–Weinberg equilibrium (HWE) was tested in females only for ChrX nonPAR (*p* ≥ 10⁻⁴); male heterozygous calls in ChrX and ChrY nonPAR were set to missing; and ChrY nonPAR sites with >10% heterozygosity across males were additionally removed as paralogous artefacts. Males in nonPAR regions were coded as pseudo-diploid homozygotes (0/0 or 1/1) prior to imputation with Beagle 5.1(Browning *et al*., 2018), using region-appropriate effective population sizes (effective population size (*ne*) = 145, 73, or 72). Post-imputation filters applied dosage *R*-squared (*DR*² ≥ 0.3), minor allele frequency (MAF) ≥ 5% for autosomes and minimum allele count (MAC) ≥ 5 for sex chromosomes, and a site-level heterozygosity filter (>50%). Pipeline verification confirmed zero heterozygous calls in male nonPAR regions, and zero female calls on ChrY. Final GWAS-ready variant counts are in Supplementary Table S4.

### 2.4 Population structure and genetic diversity

Population structure, kinship, principal components, MAF, and heterozygosity were estimated as described in Babaei and Torkamaneh (2026) using TASSEL v5 (Bradbury *et al*., 2007) and FastSTRUCTURE (Raj *et al*., 2014). Nucleotide diversity (*θπ*) and Tajima’s *D* were calculated in non-overlapping 10 kb windows using VCFtools v0.1.16 (Danecek *et al*., 2011) separately for each chromosomal compartment (PAR, HXR, PAR_Y, MSY) and sex stratum (all 145 accessions, 73 females, 72 males). Pairwise comparisons between compartments used the Wilcoxon rank-sum test with Bonferroni correction (Cuzick, 1985). The proportion of SNPs located within annotated genes was determined by intersecting variant positions with Otto II gene annotation BED files using BEDTools v2.31.1 (Quinlan and Hall, 2010; Carey *et al*., 2026); one genomic gap exceeding 1 Mb (3.3 Mb) was detected in the ChrY MSY, consistent with a centromeric region of near-zero mappability. Pairwise genetic distances were estimated with MEGA11 (Kumar et al., 2018; *p*-distance model, 1000 bootstrap replicates) per compartment and sex stratum; results are presented as a heatmap in R using the ‘*pheatmap’* package (Kolde & Kolde, 2015; Team, 2020). Linkage Disequilibrium (LD) decay was computed with PopLDdecay v3.41 (Zhang et al., 2019; maximum distance 500 kb) for each compartment and sex stratum separately. Haplotype blocks (HB) were defined with HaploView v4.2 (Barrett *et al*., 2005) using the Gabriel et al. 2002 confidence-interval method. Variant types and predicted functional impacts (intergenic, intronic, synonymous, missense) along with the transition-to-transversion (Ts/Tv) ratio were assessed using SnpEff v5.2e (Cingolani *et al*., 2012) with a custom database built from the Otto II assembly gene models (Carey *et al*., 2026). Phylogenetic trees were constructed using the Neighbour-Joining (NJ) method in MEGA11 (Kumar *et al*., 2018) under three substitution models (i.e., Jukes–Cantor (JC), Tajima–Nei (TN), and Maximum Composite Likelihood (MCL); 1000 bootstrap replicates), visualized with iTOL v7.2 (Letunic and Bork, 2021), and annotated with sample metadata comprising climate zone, biological sex, ADMIXTURE cluster, chemotype (Babaei and Torkamaneh, 2026), plant architectural type, and flowering group (Babaei *et al*., 2026*b*).

### 2.5 XY *k*-mer-based GWAS

*K*-mer-based GWAS was performed using XY_kmer_GWAS (github.com/Mehdibabaeii/XY_kmer_GWAS), a sex chromosome-aware pipeline built on the kmersGWAS framework (Voichek and Weigel, 2020), following the read-mapping and visualization approach of Lemay et al. (2023). Prior to the full analysis, four strategies varying *k*-mer count threshold (CI), strand proportion filter (PROP), kinship source, and PCR deduplication were evaluated on ChrX females using three representative traits (FT50I, SDH, TFW), comparing genomic inflation (*λ*), number of significant *k*-mers, and number of genomic signal windows (Supplementary Table S5). The final strategy (CI = 3, PROP = 0.0, *k*-mer kinship, with PCR deduplication) was selected because it applies the methodologically appropriate *k*-mer-derived kinship for each run, uses the correct strand filter for GBS data (Voichek and Weigel, 2020; Jaegle *et al*., 2025), and includes PCR deduplication and was applied to all final analyses (Supplementary Table S5).

Four independent runs were performed: ChrX females (*n* = 73), ChrX males (*n* = 72), ChrY males (*n* = 72), and whole genome (*n* = 145). Sex-stratified runs detect trait-associated *k*-mers based on within-sex variation on each sex chromosome independently, avoiding confounding by hemizygosity and sex-linked LD structure. The whole-genome run provides a complementary genome-wide view of marker–trait associations across autosomes and sex chromosomes simultaneously, capturing the contribution of sex-chromosome variation within the full genomic context.

ChrX- and ChrY-specific reads were extracted from coordinate-sorted BAM files using SAMtools v1.21 (Danecek *et al*., 2021). PCR duplicates were removed with Clumpify (BBTools v36.92; Bushnell, 2014) prior to *k*-mer counting. *K*-mer tables were constructed with KMC3 (*k* = 31, CI = 3, CS = 10,000, PROP = 0.0; Kokot et al., 2017). A strand proportion filter of 0.0 was applied because GBS libraries are inherently strand-specific, and applying a non-zero filter removes valid single-strand *k*-mers generated by restriction-site digestion (Voichek and Weigel, 2020; Jaegle *et al*., 2025). A presence/absence matrix was built retaining *k*-mers with MAC ≥ 5 across samples. Kinship matrices were computed from the full *k*-mer presence/absence table of each run independently, ensuring the covariate structure was specific to each sex-chromosome dataset following the standard kmersGWAS approach (Voichek and Weigel, 2020). Association testing first applied an approximate linear mixed model to a random sample of 1,000,001 *k*-mers to identify top candidates, followed by exact testing in GEMMA v0.96 (Zhou and Stephens, 2012) with MAF ≥ 0.05; run-specific 5% significance thresholds were established using 100 permutations.

Significant *k*-mers were mapped back to individual sample reads using katcher (Lemay et al., 2023; MQ ≥ 20), then to the Otto II reference assembly using BWA v0.7.17 (Li and Durbin, 2009). Mapped positions were clustered into signal windows within 250 kb and classified as PAR or nonPAR using the 30 Mb boundary on ChrX and ChrY. The minimum read count filter for final figures and tables was set to 10, selected based on evaluation at thresholds of 1, 5, and 10 (Supplementary Table S6). Genomic signal windows were annotated with the nearest gene model within 250 kb using the Otto II HAP2 (ChrX) and HAP1 (ChrY) primary GFF3 annotation files (Carey *et al*., 2026).

### 2.6 XY SNP-based GWAS

SNP-based GWAS for ChrX (*n* = 145) and ChrY (*n* = 72 males) was performed using the Bayesian-information and linkage-disequilibrium iteratively nested keyway (BLINK) implemented in GAPIT3 (Huang *et al*., 2019; Wang *et al*., 2022); autosomal GWAS was published previously (Babaei and Torkamaneh, 2026). Kinship matrix and principal components were derived exclusively from autosomal SNPs to prevent confounding with sex-linked signals (König *et al*., 2014) and applied as covariates in both runs; biological sex was not included as a covariate for ChrX or ChrY GWAS to avoid suppressing sex-linked associations. Genome-wide significance thresholds were set at Bonferroni-corrected *p* ≤ 0.05 divided by the number of tested SNPs per run; a suggestive threshold of *p* ≤ 1 × 10^-7^ was additionally applied. Manhattan and quantile–quantile (QQ) plots were generated within rMVP v1.0.6 (Yin *et al*., 2021) and GAPIT3. Genome-wide distribution of significant markers was visualized using shinyCircos v2.0 (Wang *et al*., 2023). Linkage disequilibrium for significant markers was characterized by HaploView v4.2 (Barrett *et al*., 2005). Functional annotation of significant variants was performed with SnpEff v5.2f using the Otto II HAP assembly database, with OIIb and OIIa primary high-confidence GFF3 annotation files for ChrX and ChrY respectively; gene function was inferred from *Arabidopsis thaliana* orthologues following Carey et al., 2026.

### 2.7 Statistical analysis

Phenotypic data summary statistics, heritability estimates, and ANOVA were reported previously (Babaei *et al*., 2024; Babaei and Torkamaneh, 2026). For the present study, descriptive statistics for genomic parameters (SNP density, MAF, heterozygosity, proportion of SNPs in genes) were calculated per chromosomal compartment and sex stratum in R v4.4 (Team, 2020). Pairwise comparisons between compartments used the Wilcoxon rank-sum test with Bonferroni correction (Cuzick, 1985). F_ST_ was calculated before and after XYRefiner filtering to validate pipeline performance; the signal-to-noise ratio was assessed as the ratio of HXR to autosomal mean F_ST_. GWAS significance thresholds are described in sections 2.5 and 2.6. Allelic effect plots for significant markers were generated with the ‘*ggplot2’* package in R (Wickham *et al*., 2016; Team, 2020)

## 3. RESULTS

### 3.1 Sex chromosome characterization

XYRefiner filtering substantially improved differentiation between sex chromosome compartments and the genomic background. After filtering, mean autosomal F_ST_ decreased to 0.003–0.004, whereas HXR F_ST_ increased to 0.108, with 86.2% of windows exceeding the genome-wide 95^th^ percentile. This resulted in a 27-fold increase in signal-to-noise ratio. In addition, no female genotypes remained on ChrY after filtering, confirming effectiveness of the sex-aware filtering approach (Supplementary Fig. S3).

The GWAS-ready variant set comprised 20,646 SNPs on ChrX (243.7 SNPs/Mb) and 6,217 SNPs on ChrY (53.0 SNPs/Mb). Compartment-level analysis revealed pronounced differences in genomic parameters (Supplementary Table S7). ChrX HXR showed the highest SNP density (283 SNPs/Mb), while ChrY MSY was markedly depleted (18.3 SNPs/Mb), consistent with its repeat-rich, gene-poor composition. The proportion of SNPs within annotated genes was highest in PAR regions (ChrX PAR: 19.8%; ChrY PAR: 17.2%) and near-absent in the MSY (0.6%), reflecting progressive degeneration of the male-specific region. Heterozygosity in males was zero across both ChrX HXR and ChrY MSY, confirming hemizygosity in these compartments. One genomic gap of 3.3 Mb was identified in the ChrY MSY, consistent with a centromeric region of near-zero mappability. The majority of annotated variant impacts were non-coding (MODIFIER class: ChrX 97.3%, ChrY 96.6% of all annotations); among coding variants, the missense-to-silent ratio was higher on ChrX (1.35) than ChrY (0.98), and the Ts/Tv ratio was lower on ChrY (1.73) than ChrX (1.92), both consistent with reduced purifying selection and mutation accumulation in the non-recombining MSY.

### 3.2 Population structure and evolutionary differentiation of sex chromosomes

ChrX phylogenetic analysis resolved males and females into two fully distinct clades (Fig. 1A). Within the female clade, two geographically and phenotypically divergent lineages were identified: Ancestry 1 comprised warm/hot-adapted accessions (BWh, BSh, Cfa) with THC-dominant chemotype and a mixture of early and late-flowering types; Ancestry 2 was exclusively cold/arid (BSk, BWk) with CBD-dominant chemotype, PT4 (menorah) plant architecture, and a combination of late-flowering and auto-flowering accessions, reflecting strong climate–chemotype–morphology co-structure on ChrX. ChrY analysis resolved two male lineages (*K* = 2; Fig. 1B); auto-flowering accessions with PT1 (lateral-attached) plant architecture appeared at intermediate positions within both lineages, predominantly from cold regions, suggesting these accessions may represent an ancestral or admixed component of the male gene pool. Pairwise genetic distances confirmed compartment-dependent differentiation across all sex strata (Supplementary Fig. S4). Within-group distances were lowest in PAR regions for both ChrX and ChrY, reflecting shared recombination history in this pseudoautosomal compartment. The ChrY MSY showed the highest within-group distances, consistent with mutation accumulation in the absence of recombination.

**Fig. 1.**
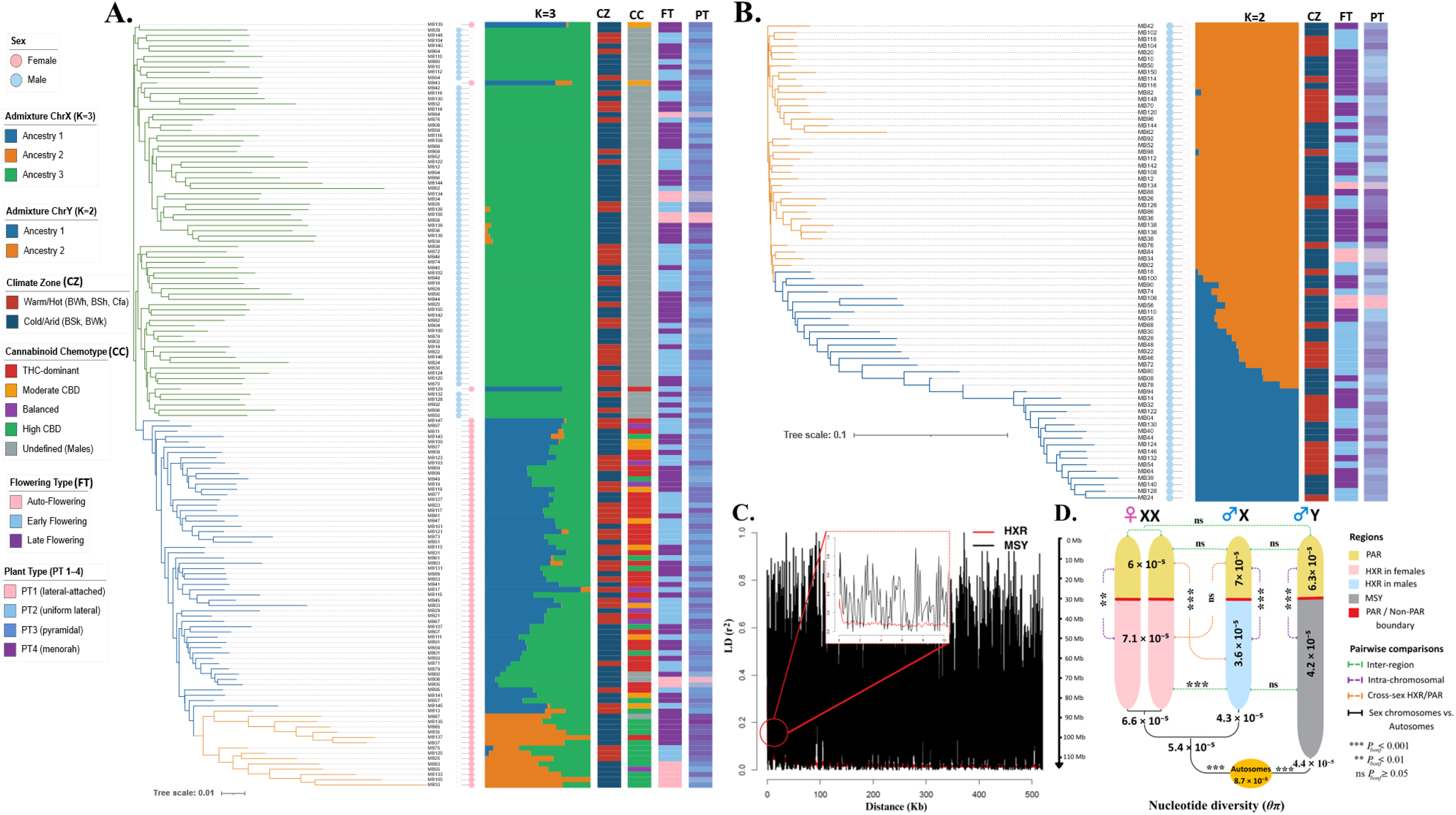
Population structure, linkage disequilibrium, and nucleotide diversity across cannabis sex chromosomes. (**A**) Neighbour-Joining phylogeny of study accessions based on ChrX variants, annotated with biological sex, ChrX admixture proportions (*K* = 3), climate zone, cannabinoid chemotype, flowering type, and plant architectural type. (**B**) Neighbour-Joining phylogeny based on ChrY variants with corresponding admixture bar plots (*K* = 2), revealing two distinct male-lineage ancestries. (**C**) Linkage disequilibrium (*r*²) decay as a function of inter-marker distance for the hemizygous X region (HXR; red) and male-specific Y region (MSY; black), with inset highlighting 0–500 kb range. (**D**) Nucleotide diversity (*θ*π) across PAR, HXR, and MSY compartments for female (XX), male X, and male Y chromosomes compared with autosomal diversity; pairwise significance levels indicated (*** *p* ≤ 0.001; ** *p* ≤ 0.01; ns *p* ≥ 0.05). Alt text: Four-panel figure. A: Neighbour-joining phylogenetic tree based on ChrX variants annotated with biological sex, ChrX admixture proportions, climate zone, cannabinoid chemotype, flowering type, and plant architectural type. B: Neighbour-joining phylogenetic tree based on ChrY variants with admixture bar plots showing two distinct male-lineage ancestries. C: Line graph of linkage disequilibrium r-squared decay against inter-marker distance for HXR in red and MSY in black with inset highlighting 0 to 500 kb range. D: Figure showing nucleotide diversity across PAR, HXR, and MSY compartments for female, male X, and male Y chromosomes compared with autosomal diversity with pairwise significance indicators.

Linkage disequilibrium (LD) patterns were strikingly compartment-dependent (Table S6; Fig. 1C; Supplementary Fig. S5). In PAR regions, *r*² dropped to half its maximum within 200–400 bp, comparable to autosomal background. The ChrY MSY showed dramatically elevated and persistent LD, with *r*² half-decay at 77,600 bp and maximum *r*² of 0.667, consistent with complete absence of recombination. Haplotype blocks in the MSY were correspondingly large (mean span 186 kb; maximum 499 kb) relative to PAR regions (mean span 0.4–0.8 kb). Sex-stratified analysis of ChrX revealed that HXR females harboured substantially more haplotype blocks (1,513; 26.5/Mb) than HXR males (290; 5.2/Mb), reflecting the diploid versus hemizygous state of this compartment.

Nucleotide diversity (*θ*π) differed significantly between compartments and sexes (Fig. 1D; Supplementary Fig. S6). All PAR pairwise comparisons were non-significant (ChrX PAR females: 6 × 10⁻⁵; males: 7 × 10⁻⁵; ChrY PAR: 6.3 × 10⁻⁵; *p* = ns for all), confirming free recombination across pseudoautosomal regions. In the HXR, females showed significantly higher *θ*π (7.1 × 10⁻⁵) than males (3.60 × 10⁻⁵; *p* < 0.001), reflecting hemizygous inheritance. HXR males were significantly lower than PAR males (*p* < 0.001), providing independent validation of the 30 Mb PAR/nonPAR boundary. HXR males and MSY showed comparable diversity (3.60 vs 4.20 × 10⁻⁵; *p* = ns), suggesting similar evolutionary constraint across both hemizygous compartments. Male ChrX and ChrY were not significantly different (4.3 vs 4.4 × 10⁻⁵; *p* = ns). All sex chromosome compartments showed significantly reduced *θ*π relative to autosomes (8.7 × 10⁻⁵; *p* < 0.001). Tajima’s *D* was consistently negative across PAR compartments and positive in the MSY, consistent with relaxed purifying selection in the non-recombining region (Supplementary Fig. S6).

### 3.3 XY-linked loci for agronomic traits

*K*-mer tables comprised 6.74M, 5.87M, 7.41M, and 100.3M *k*-mers for ChrX_females, ChrX_males, ChrY_males, and whole_genome runs respectively. Sex-stratified runs identified 506, 275, and 136 signal windows for ChrX_females (426 nonPAR; 80 PAR), ChrX_males (187 nonPAR; 88 PAR), and ChrY_males (117 nonPAR; 19 PAR) respectively; the whole-genome run yielded 2,572 signals across ChrY (667), ChrX (254), and autosomes (1,651; Supplementary Table S8; Supplementary Figs S7, S8). Fifteen traits yielded concordant signals across all four runs — DWS, FT50I, FWS, HGV, HH, LLLS, LMI, NMI, NNH, NTFIS, NTMI, RGR, SDH, SFFI, and TDW — identifying these as robustly sex-chromosome-associated loci (Supplementary Table S8). The strongest signals in sex-stratified runs were RGR at 50.8 Mb in ChrX_females (−log_10_ (*p*)= 17.7), SDH at 58.6 Mb in ChrX_males (−log_10_ (*p*)= 18.2), and RGR at 106.4 Mb in ChrY_males (−log_10_ (*p*)= 15.2), with genomic coordinates, flanking gene annotations, and nearest SNP markers for all 1,838 signal windows provided in Supplementary Table S9. Across sex-stratified runs, 262, 162, and 102 unique genomic windows were resolved for ChrX_females, ChrX_males, and ChrY_males respectively; the whole-genome run identified 100 unique ChrX windows and 456 ChrY windows (Supplementary Table S8). The most pleiotropic hotspot was at 42.1 Mb on ChrX (18 traits; −log_10_ (*p*) = 24.4), followed by 58.6 Mb on ChrX (17 traits; −log_10_ (*p*) = 18.2) and 106.1 Mb on ChrY (10 traits; −log_10_ (*p*) = 17.7; Supplementary Fig. S9).

Sex-stratified signal patterns revealed pronounced asymmetries between compartments and sexes (Supplementary Table S10). In the HXR, biomass traits were strongly female-biased on ChrX (TFW: 55 female vs 1 male windows; TDW: 22 vs 1; SDH: 43 vs 12), while DWF, RGR, and NLS were exclusively male-detected. GVP, LIMTH, and LSLS yielded signals only in females on ChrX nonPAR. In contrast, 13 traits showed concordant signals in both sexes in the PAR — FT50I, FWS, HH, LLLS, LMI, NMI, NNH, NTFIS, RGR, SDH, SF10I, SF10P, and SFFI — consistent with pseudoautosomal inheritance. In the whole-genome run, biomass traits showed strong enrichment on ChrY across the MSY region (TDW: 107 signals; DWS: 101; TFW: 97; FT50I: 88). LD analysis revealed a highly complex, trait-specific genetic architecture across the sex chromosomes. While ChrY_males signals generally resolved to fewer independent loci (median N_independent_loci = 1.5), they also harbored notable exceptions, such as RGR resolving to 13 independent loci. Moreover, both the male and female X chromosomes exhibited striking structural complexity. In ChrX_females, traits like LMI, RGR, and SDH each resolved to 11 independent loci. Interestingly, the male X chromosome demonstrated an equally pronounced, if not greater, architectural complexity. This was most evident for RGR, which peaked at 21 independent loci on ChrX_males, alongside highly complex architectures for NNH (11 loci) and SDH (8 loci). This suggests that structural complexity is strongly trait-driven rather than solely a product of female X chromosome diploidy (Supplementary Table S11). 22 *k*-mer signal windows co-localized with significant SNP loci, providing independent cross-validation for eight ChrX and three ChrY markers (Supplementary Table S12).

SNP-based GWAS on ChrX (*n* = 145) identified eight significant genome-wide markers across six agronomic traits, with Phenotypic Variance Explained (PVE) values ranging from 4.4% (CsX5 for FWF) to 73.6% (CsX8 for RGR; Table 1; Fig. 2A; Supplementary Figs S10A, S11, S13). Quantile–quantile plots confirmed appropriate model fit with no global inflation of test statistics (Supplementary Fig. S10A). Seven markers were located in the HXR and one in the PAR. The pleiotropic locus *StemWeightLocus1* (*CsWSL1*; CsX1; ChrX: 59,815,662) was significantly associated with both DWS (*p* = 3.1 × 10^-9^; PVE = 56.4%) and FWS (*p* = 1.9 × 10^-8^; PVE = 51%), with the major “A” allele reducing DWS by 17.6 g and FWS by 27.9 g. FWF was controlled by four HXR markers (CsX2–CsX5; 42.5–74.5 Mb; PVE 4.4–32.9%): the major “A” allele of CsX3 (ChrX: 48,968,807; *p* = 3.5 × 10^-15^) increased FWF by 49.7 g (PVE = 32.9%), the largest single-marker effect among FWF loci, while the major “T” allele of CsX5 (ChrX: 74,517,203) reduced FWF by 13.7 g (PVE = 4.4%), suggesting antagonistic regulation across the HXR. CsX7 (ChrX: 41,340,487) explained 63% of NTMI variance (*p* = 6.9 × 10^-8^), and CsX8 (ChrX: 52,199,006) showed the highest single-marker PVE on ChrX (73.6% for RGR; *p* = 1.6 × 10^-10^). The sole PAR marker, CsX6 (ChrX: 14,515,776), was significantly associated with LLLS (*p* = 8.9 × 10^-11^; PVE = 39.7%), with the major “T” allele increasing lateral shoot length by 20.9 cm, consistent with pseudoautosomal inheritance and concordant *k*-mer signals in both sexes (windows kXF014–016, kXM012; Supplementary Table S12). Haplotype block analysis confirmed defined LD structure at seven of the eight ChrX markers; no block was detected at CsX3 (Supplementary Fig. S13; Supplementary Table S12). Block spans ranged from 0.03 kb (CsX2) to 7.9 kb (CsX4), reflecting elevated recombination in the HXR.

**Fig. 2.**
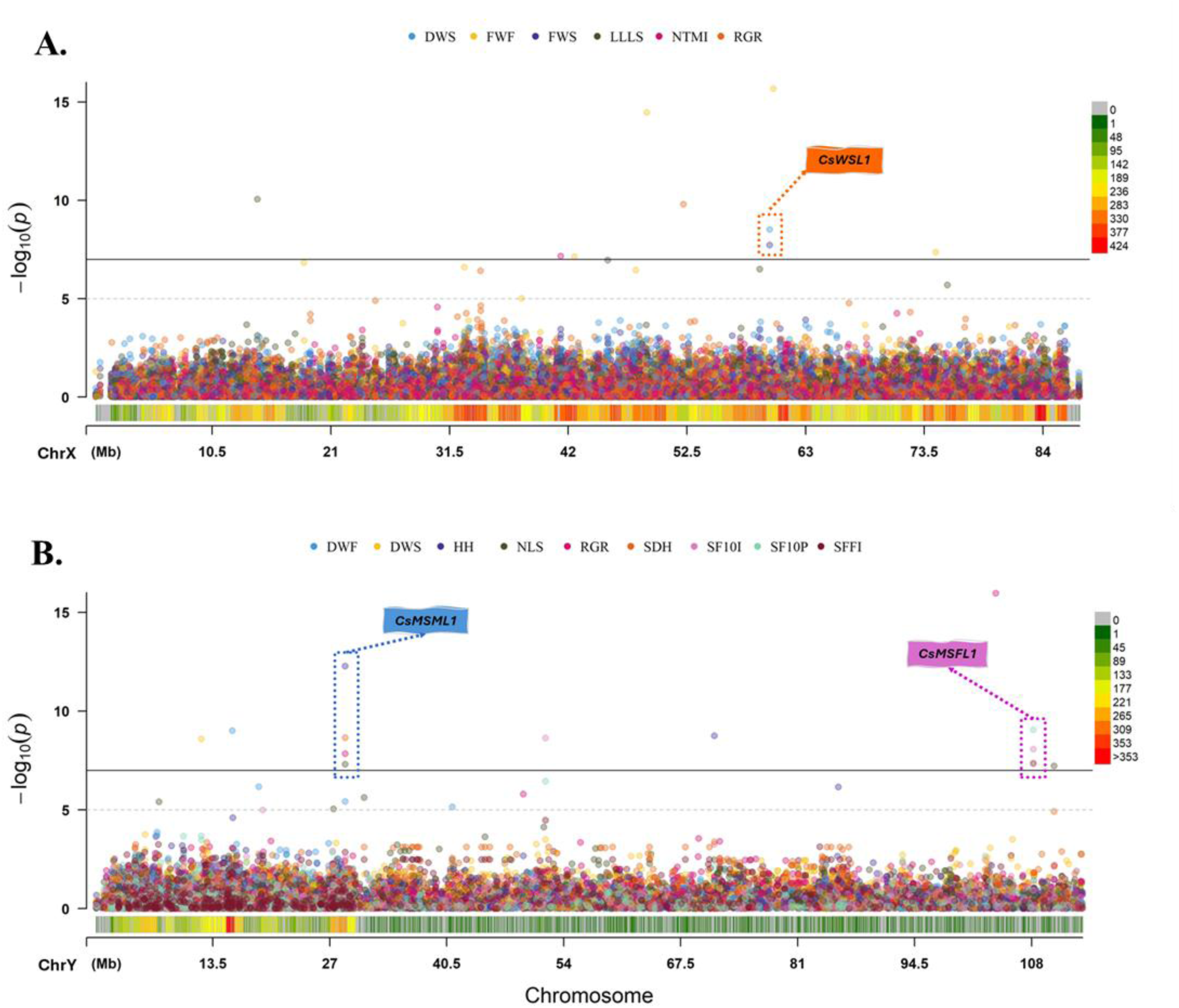
Manhattan plots of SNP-based GWAS signals on cannabis sex chromosomes. Association signals (−log10 (*p*)) plotted across (**A**) ChrX (n = 145) and (**B**) ChrY (n = 72 males) for 27 agronomic traits. The colour bar below each plot represents local SNP density. Genomic positions below and above 30 Mb correspond to the pseudoautosomal region (PAR) and non-recombining compartments (HXR on ChrX; MSY on ChrY), respectively. Notable loci are indicated: *CsWSL1* (ChrX, 59.8 Mb), *CsMSML1* (ChrY, 28.8 Mb) and *CsMSFL1* (ChrY, 108.2 Mb). Alt text: Two-panel Manhattan plot. A: Association signals plotted across ChrX for 145 accessions and 27 agronomic traits with SNP density color bar and CsWSL1 locus indicated at 59.8 Mb. B: Association signals plotted across ChrY for 72 males and 27 agronomic traits with SNP density color bar and CsMSML1 at 28.8 Mb and CsMSFL1 at 108.2 Mb indicated.

**Table 1.** Genome-wide significant SNP markers associated with agronomic traits on cannabis ChrX and ChrY.

| Traits | Marker ID | Locus ID<sup>a</sup> | Chr | Position (bp)<sup>b</sup> | Compartment<sup>c</sup> | *p*-value | MAF<sup>d</sup> (%) | Allele<sup>e</sup> | Effect<sup>f</sup> | PVE<sup>g</sup> (%) |
| --- | --- | --- | --- | --- | --- | --- | --- | --- | --- | --- |
| DWS | CsX1 | *CsWSL1* | X | 59815662 | HXR | 3.06E-09 | 8.3 | A/G | -17.57 | 56.43 |
| FWF | CsX2 | - | X | 42549044 | HXR | 7.38E-08 | 4.8 | C/T | 22.157 | 8.12 |
|  | CsX3 | - | X | 48968807 | HXR | 3.45E-15 | 2.1 | A/C | 49.692 | 32.9 |
|  | CsX4 | - | X | 60151058 | HXR | 2.12E-16 | 5.5 | G/C | 32.242 | 29.78 |
|  | CsX5 | - | X | 74517203 | HXR | 4.28E-08 | 15.2 | T/C | -13.664 | 4.43 |
| FWS | CsX1 | *CsWSL1* | X | 59815662 | HXR | 1.88E-08 | 8.3 | A/G | -27.881 | 50.96 |
| LLLS | CsX6 | - | X | 14515776 | PAR | 8.89E-11 | 3.4 | T/A | 20.863 | 39.71 |
| NTMI | CsX7 | - | X | 41340487 | HXR | 6.89E-08 | 4.1 | T/C | 2.268 | 63.02 |
| RGR | CsX8 | - | X | 52199006 | HXR | 1.6E-10 | 1.7 | C/T | -11.529 | 73.62 |
| DWF | CsY1 | - | Y | 15770838 | PAR | 1E-09 | 48.6 | A/G | 1.872 | 8.55 |
| DWS | CsY2 | - | Y | 12186985 | PAR | 2.62E-09 | 8.3 | A/T | -16 | 69.29 |
| HH | CsY3 | *CsMSML1* | Y | 28768718 | PAR | 5.37E-13 | 3.5 | G/T | 68.223 | 35.8 |
|  | CsY4 | - | Y | 71393088 | MSY | 1.78E-09 | 2.8 | T/A | 36.33 | 45.1 |
| NLS | CsY3 | *CsMSML1* | Y | 28768718 | PAR | 5.03E-08 | 3.5 | G/T | 5.878 | 32.15 |
|  | CsY5 | - | Y | 110611039 | MSY | 6.03E-08 | 33.3 | T/C | 2.898 | 15.07 |
| RGR | CsY3 | *CsMSML1* | Y | 28768718 | PAR | 1.47E-08 | 3.5 | G/T | -14.509 | 17.18 |
|  | CsY6 | - | Y | 103862024 | MSY | 1.1E-16 | 4.2 | C/T | -17.813 | 51.25 |
| SDH | CsY3 | *CsMSML1* | Y | 28768718 | PAR | 2.3E-09 | 3.5 | G/T | 4.232 | 76.13 |
| SF10I | CsY7 | - | Y | 51928260 | MSY | 2.32E-09 | 44.4 | A/C | -8.192 | 8.26 |
|  | CsY8 | *CsMSFL1* | Y | 108197200 | MSY | 8.58E-09 | 6.9 | A/C | -19.441 | 84.9 |
| SF10P | CsY8 | *CsMSFL1* | Y | 108197200 | MSY | 8.97E-10 | 6.9 | A/C | -24.897 | 84.83 |
| SFFI | CsY8 | *CsMSFL1* | Y | 108197200 | MSY | 4.51E-08 | 6.9 | A/C | -15.617 | 87.08 |

<sup>a</sup> Complete name of key introduced loci: *StemWeightLocus1* (*CsWSL1*); *Male-SpecificMorphologyLocus1* (*CsMSML1*); *Male-SpecificFloweringLocus1* (*CsMSFL1*).
<sup>b</sup> Genomic positions reported based on the Otto II haplotype-resolved assembly (OIla haplotype for ChrY; OIIb haplotype for ChrX).
<sup>c</sup> Chromosomal compartments: PAR = Pseudoautosomal Region (shared by X and Y, position < 30 Mb); HXR = Hemizygous X Region (ChrX nonPAR, position > 30 Mb); MSY = Male-Specific Y Region (ChrY nonPAR, position > 30 Mb).
<sup>d</sup> Minor Allele Frequency.
<sup>e</sup> Major/minor.
<sup>f</sup> Estimated allelic effect of the major allele on the trait.
<sup>g</sup> Proportion of Phenotypic Variance Explained by the significant SNP.

SNP-based GWAS on ChrY (*n* = 72 males) identified eight genome-wide significant markers across nine agronomic traits spanning PAR (*n* = 3) and MSY (*n* = 5), with PVE values ranging from 8.3% (CsY7 for SF10I) to 87.1% (CsY8 for SFFI; Table 1; Fig. 2B; Supplementary Figs S10B, S12, S14). Quantile–quantile plots confirmed appropriate model calibration for the male-only analysis (Supplementary Fig. S10B). The pleiotropic locus *Male-SpecificMorphologyLocus1* (*CsMSML1*; CsY3; ChrY: 28768718 PAR) was associated with HH (*p* = 5.4 × 10^-13^; PVE = 35.8%), SDH (*p* = 2.3 × 10^-9^; PVE = 76.1%), RGR (*p* = 1.5 × 10^-8^; PVE = 17.2%), and NLS (*p* = 5 × 10^-8^; PVE = 32.2%), where the major “G” allele increased plant height by 68.2 cm and stem diameter by 4.2 mm. CsY2 (ChrY: 12,186,985 PAR) explained 69.3% of DWS variance (*p* = 2.6 × 10^-9^), with the major “A” allele reducing stem dry weight by 16 g. The most statistically significant ChrY marker was CsY6 (ChrY: 103,862,024 MSY; *p* = 1.1 × 10^-16^; PVE = 51.3% for RGR), where the major “C” allele reduced relative growth rate by 17.8 units, co-localizing with *k*-mer window kYM092 (Supplementary Table S12). *Male-SpecificFloweringLocus1* (*CsMSFL1*; CsY8; ChrY 108,197,200 MSY) was associated with SFFI (*p* = 4.5 × 10^-8^; PVE = 87.1%), SF10P (*p* = 9 × 10^-10^; PVE = 84.8%), and SF10I (*p* = 8.6 × 10^-9^; PVE = 84.9%), where the major “A” allele delayed flowering by 15.6, 24.9, and 19.4 days respectively — the highest PVE values observed across the entire dataset. Haplotype block analysis revealed markedly extended LD at MSY markers relative to PAR (Supplementary Fig. S14; Supplementary Table S12). MSY block spans ranged from 136 kb (CsY4) to 493 kb (CsY5), consistent with suppressed recombination in the non-recombining region, while PAR blocks were substantially smaller (CsY3: 0.04 kb; CsY1: 0.01 kb), with the exception of CsY2, which showed an unusually extended block of 110 kb suggesting locally reduced recombination in the PAR.

Candidate gene analysis within haplotype block regions flanking the 16 significant markers identified putative regulatory functions consistent with their associated traits (Supplementary Table S12; Fig. 3). *CsWSL1* (CsX1; ChrX:59,815,662) flanks a MATE-family membrane transporter (*OIIb.chrX.v1.g424870*) and a RING-H2 ubiquitin ligase (*OIIb.chrX.v1.g424880*), implicating membrane-mediated solute transport and targeted protein degradation in stem biomass accumulation. *CsMSML1* (CsY3; ChrY:28,768,718) is flanked bilaterally by a G-type lectin S-receptor-like serine/threonine kinase (*OIIa.chrY.v1.g384890*), consistent with a role in cell-to-cell signaling underlying the coordinated control of height, diameter, and branching. Notably, CsX3 (ChrX:48,968,807) and CsY6 (ChrY:103,862,024) each flank a ribonuclease H-like protein on independent sex chromosomes (*OIIb.chrX.v1.g420950* and *OIIa.chrY.v1.g399450* respectively), suggesting a conserved RNA-processing function at convergent loci. *CsMSFL1* (CsY8; ChrY:108,197,200) flanks currently uncharacterized genes; its exclusive MSY location, 235 kb haplotype block and high PVE across three phenological traits support a male-specific regulatory function in flowering time control. The genome-wide co-distribution of all significant SNP and *k*-mer markers, revealing spatial co-clustering between 40–60 Mb on ChrX and 100–110 Mb on ChrY, is presented in a circular genomic overview (Fig. 3).

**Fig. 3.**
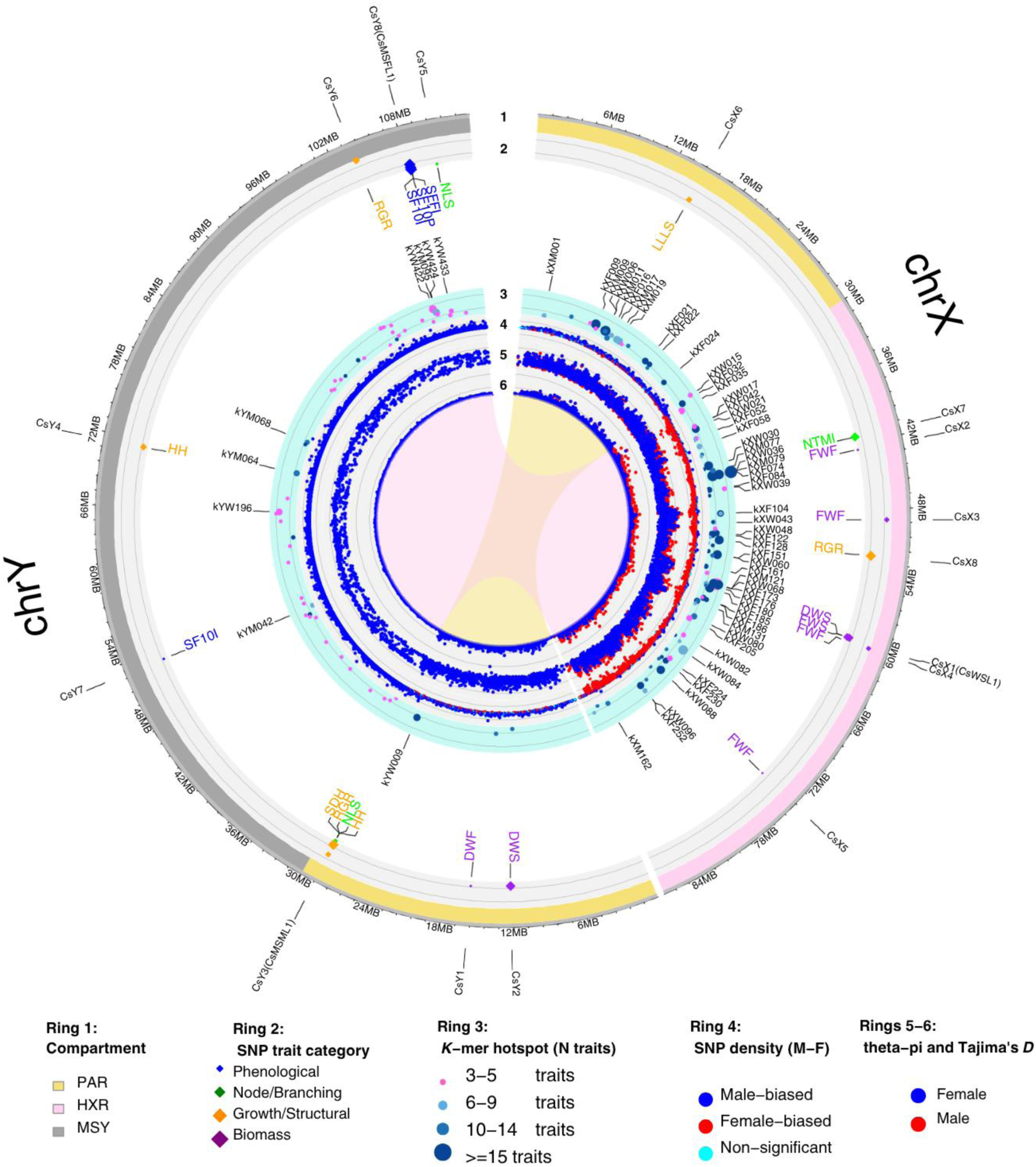
Circular genomic overview of cannabis sex chromosomes. The outer ring shows chromosomal compartments for ChrX and ChrY: pseudoautosomal region (PAR; yellow), hemizygous X region (HXR; pink), and male-specific Y region (MSY; gray). Moving inward: (Ring 2) genome-wide significant SNP-GWAS markers, colored by trait subcategory; (Ring 3) pleiotropic *k*-mer GWAS hotspot windows, with color intensity reflecting the number of associated traits and point size scaled accordingly; (Ring 4) SNP density sex-differentiation index; (Ring 5) windowed nucleotide diversity (*θ*π) and (Ring 5) Tajima’s *D*, with female accessions in red and male accessions in blue. Alt text: Circular genomic figure with concentric rings for ChrX and ChrY. Outer ring shows PAR in yellow, HXR in pink, and MSY in gray. Inner rings show genome-wide significant SNP-GWAS markers colored by trait subcategory, pleiotropic k-mer hotspot windows with color intensity and point size scaled by number of associated traits, SNP density sex-differentiation index, windowed nucleotide diversity and Tajima's D with female accessions in red and male accessions in blue.

## 4. DISCUSSION

This study presents the first sex chromosome-aware GWAS of *Cannabis sativa*, integrating both ChrX and ChrY into a unified analytical framework across 145 landrace accessions and 27 agronomic traits. By combining SNP-based GWAS with sex-stratified *k*-mer association analysis, and leveraging the Otto II haplotype-resolved assembly (Carey *et al*., 2026), we identified 16 genome-wide significant SNP markers across both sex chromosomes: eight on ChrX controlling six agronomic traits including stem biomass (*CsWSL1*), lateral shoot architecture (CsX6), and relative growth rate (CsX8), and eight on ChrY controlling nine traits including plant morphology (*CsMSML1*) and male-specific flowering (*CsMSFL1*). In parallel *k*-mer GWAS yielded 1,838 annotated signal-trait associations representing 1,082 unique genomic windows distributed across PAR, HXR, and MSY compartments. This scale and resolution were unattainable in previous cannabis GWAS studies where Y chromosome sequences were absent (de Ronne *et al*., 2024; de Ronne and Torkamaneh, 2025; Babaei and Torkamaneh, 2026). Beyond their evolutionary significance, the 16 sex-linked loci identified here constitute the first genomic markers directly applicable to paternal line selection and sex-chromosome-aware crop improvement in cannabis. The three chromosomal compartments — PAR, HXR, and MSY — confer fundamentally different inheritance modes that determine how each locus can be deployed in breeding. PAR markers such as *CsMSML1* behave as conventional co-dominant Mendelian loci transmissible through either parent, directly applicable to standard marker-assisted selection in both males and females. HXR markers including *CsWSL1* and CsX8 are hemizygous in males and diploid in females; selecting through female lines can expose recessive alleles masked by hemizygosity — a property unique to X-linked loci and exploitable in backcross or recurrent selection schemes. MSY markers, particularly *CsMSFL1*, are strictly paternally inherited: a single Y-linked genotype call at the seedling stage can predict male flowering phenology with near-deterministic accuracy, enabling paternal curation before plants reach anthesis and before any field resources are committed to maintaining suboptimal males. The broader significance of these results lies in what they reveal about a systematic blind spot in cannabis genomics: male plants, long treated as an inconvenience to be rogued or a mere pollen source, carry a Y-chromosomal and X-linked genetic architecture of substantial agronomic consequence (Welling *et al*., 2020; Lynch *et al*., 2025; Wei *et al*., 2026). In dioecious plants and animals alike, the genetic architecture of the male sex is rarely investigated with the same rigour as the female (Muyle *et al*., 2021; Sun *et al*., 2023; Charlesworth and Harkess, 2024).

The sex chromosome structure revealed by our population-level analysis tells a coherent evolutionary story consistent with the ancient history of recombination suppression in *Cannabis* and *Humulus*, estimated at approximately 28–48 Mya depending on the method and markers applied (Prentout *et al*., 2021; Carey *et al*., 2026). The PAR/nonPAR boundary at 30 Mb, determined by sex-differentiated read coverage analysis and SNP density permutation testing, and corroborated by downstream LD half-decay distances, *θ*π discontinuity, trait signal compartmentalization, and pseudoautosomal marker inheritance, aligns closely with the assembly-based boundary of 29.3 Mb reported by Carey et al., 2026, providing the first population-genetic confirmation of this structural feature from a GWAS panel. The nucleotide diversity patterns across compartments reflect three biologically distinct regimes. In the PAR, *θ*π was statistically indistinguishable among females (6 × 10^-5^), males (7 × 10^-5^), and ChrY (6.3 × 10^-5^), confirming unrestricted recombination homogenizes diversity between the sexes in this compartment. In the HXR, female *θ*π (7.1 × 10^-5^) significantly exceeded male *θ*π (3.6 × 10^-5^; *p* < 0.001), a direct consequence of hemizygosity: the effective population size for X-linked variants in hemizygous males is half that in diploid females, reducing diversity accordingly (Muyle *et al*., 2021). The absence of a significant difference between HXR males and the MSY (3.6 vs. 4.2 × 10^-5^; *p* = ns) is consistent with Muller’s ratchet operating in the non-recombining region: the accumulation of slightly deleterious mutations under relaxed purifying selection maintains the number of segregating sites even as mutation quality deteriorates, a pattern corroborated by the lower Ts/Tv (1.73 vs. 1.92) and missense-to-silent ratio (0.98 vs. 1.35) on ChrY relative to ChrX (Charlesworth, 2021). All sex chromosome compartments showed significantly reduced *θ*π relative to autosomes (8.7 × 10^-5^; *p* < 0.001), consistent with the theoretically expected reduction in effective population size for sex-linked regions under male heterogamety. ChrY phylogenetic analysis resolved two deeply divergent male lineages (*K* = 2), consistent with the half-sibling family structure characteristic of wind-pollinated dioecious plants, wherein a single female may be fertilized by pollen from multiple genetically distinct males (Friedman and Barrett, 2011). Under this reproductive system, Y chromosome diversity is shaped not only by drift and mutation but also by the selective advantage of males whose flowering time coincides with female receptivity (Sandler *et al*., 2018; Delph, 2019), suggesting that the divergence between the two ChrY lineages may partly reflect differential selection on male phenology across the Iranian landrace populations sampled here. The two divergent ChrY lineages identified here echo the two MSY haplogroups resolved in Bactrian camels, one wild and one domestic, where paternal introgression was detected in a wild individual carrying a domestic Y haplotype (Felkel *et al*., 2019); the intermediate positioning of auto-flowering accessions between both cannabis ChrY lineages raises an analogous possibility of historical male-mediated gene flow between otherwise divergent landrace groups.

Perhaps the most striking pattern to emerge from the *k*-mer analysis is the profound asymmetry in signal architecture between the sexes, an asymmetry that reflects fundamental differences in developmental investment. Female plants invest heavily in sustained inflorescence biomass over extended reproductive periods (mean TFW: 278 vs 151 g; FWF: 88 vs 29 g in females vs males); male plants prioritize rapid vegetative growth and early pollen dispersal before senescing, exhibiting significantly higher relative growth rates (RGR: 0.074 vs 0.069 g g⁻¹ day⁻¹; Babaei et al., 2024), a life-history strategy that may be partly encoded on the sex chromosomes themselves. This biology is written directly into the *k*-mer signal landscape: TFW yielded 55 female windows versus one male window on ChrX nonPAR, TDW 22 versus one, and SDH 43 versus 12, while DWF and NLS were exclusively male-detected and RGR, with 80 male versus 41 female nonPAR windows and 25 male versus 9 female PAR windows on ChrX, and the highest signal count of any trait on ChrY (69 windows), was predominantly male-associated across all sex chromosome compartments, consistent with the higher relative growth rates documented in male plants (Babaei and Ajdanian, 2020; Babaei *et al*., 2022, 2024, 2026*b*). This sex-specific asymmetry echoes a fundamental property of dioecious genomes: Torres et al. (2018) demonstrated in date palm that 1,653 *k*-mers were conserved across all 13 *Phoenix* males yet no *k*-mer was conserved exclusively across all females, consistent with male-biased sequence enrichment in the MSY of XY systems. The pleiotropic hotspots at 42.1 Mb (18 traits; −log_10_ (*p*)= 24.4) and 58.6 Mb (17 traits) on ChrX, and 106.1 Mb on ChrY (10 traits; −log_10_ (*p*)= 17.7), suggest regulatory hubs coordinating broad developmental programs across the HXR and MSY, consistent with co-option of sex chromosomes for quantitative trait regulation (Rice, 1984). The 22 windows co-localizing between *k*-mer and SNP GWAS provide independent cross-validation, and the 456 unique MSY windows from the whole-genome run alone demonstrate that the cannabis Y chromosome retains a genomically active agronomic landscape inaccessible to any GWAS design lacking a phased Y assembly or male-only samples.

The candidate genes flanking the most significant SNP markers illuminate biologically coherent molecular mechanisms. *CsWSL1* (ChrX:59,815,662; PVE 56.4% for stem dry weight and 51% for stem fresh weight) flanks a MATE-family plasma membrane organic acid efflux transporter and a RING-H2 ubiquitin ligase. MATE transporters mediate organic solute efflux across plant membranes and have been implicated in seedling development and hormone signaling (Magalhaes *et al*., 2007; Wang *et al*., 2017), while RING-H2 E3 ligases constitute a conserved family with roles in plant growth, development, and stress-responsive protein turnover (Zhang *et al*., 2007; Kong *et al*., 2025), together implicating membrane-mediated transport and targeted protein degradation in stem biomass regulation at this X-linked locus. *CsMSML1* (ChrY:28,768,718, PAR; PVE up to 76.1% for stem diameter, 35.8% for plant height, 32.2% for lateral shoot number, and 17.2% for relative growth rate) is flanked bilaterally by a G-type lectin serine/threonine receptor kinase. Lectin receptor-like kinases are established regulators of developmental pathways, stress responses, and hormonal signaling in plants (Vaid *et al*., 2013). Despite its pseudoautosomal location and presence on both X and Y chromosomes, *CsMSML1* yielded significant associations exclusively in the male GWAS panel, suggesting that its pleiotropic control of male plant morphology, spanning stem dimensions, branching architecture, and growth rate simultaneously, operates through a sex-specific regulatory context not detectable in the mixed-sex ChrX analysis, possibly reflecting sex-differential expression or epistatic interactions specific to the XY genetic background. The convergence of CsX3 (ChrX:48,968,807) and CsY6 (ChrY:103,862,024) on ribonuclease H-like flanking genes on independent sex chromosomes is particularly notable: RNase H enzymes resolve RNA:DNA hybrids that arise during replication and repair, with failure to do so leading to genome instability in higher eukaryotes (Cerritelli and Crouch, 2009), and their independent appearance at GWAS peaks on both sex chromosomes raises the possibility that RNA:DNA hybrid resolution contributes to sex-linked trait variation in cannabis. One of the most consequential for applied genomics is *CsMSFL1* (ChrY:108,197,200, MSY; PVE 84.8–87.1% for male flowering initiation), located within a 234.8 kb MSY haplotype block flanked by currently uncharacterized genes. Male cannabis produces open, highly branched staminate inflorescences adapted for prolific anemophilous pollen dispersal (Lavie *et al*., 2026), in contrast to the compact, trichome-bearing pistillate inflorescences of females optimized for pollen capture (Spitzer-Rimon *et al*., 2019). From an evolutionary perspective, males that flower before female receptivity peaks senesce before achieving maximum pollen transfer, reducing their probability of Y chromosome transmission to subsequent generations, a reproductive fitness cost that may have driven selection on *CsMSFL1* alleles that synchronize male flowering with female anthesis (Matsuhisa and Ushimaru, 2015). Compared with the autosomal flowering loci identified in our previous study, *CsFTL3*, *CsFTL4*, and *CsCFL1*, controlling phenological traits with PVE up to 44.7% across the mixed-sex panel (Babaei and Torkamaneh, 2026; Babaei *et al*., 2026*b*), *CsMSFL1* exceeds these effects substantially and acts exclusively within the male developmental program, underscoring the qualitative distinction between autosomal and sex-linked flowering regulation in this species. The applied breeding implications of *CsMSFL1* are direct and substantial. In controlled crossing programs and open-pollinated field nurseries, synchronization of male pollen shed with female receptivity is a primary determinant of seed set: males flowering too early senesces before peak female receptivity, while late-flowering males miss the optimal fertilization window entirely (Matsuhisa and Ushimaru, 2015; Lipson Feder *et al*., 2021). With *CsMSFL1* accounting for 84–87% of variance in three phenological traits, seedling-stage genotyping could replace the current practice of growing males to full anthesis before assessing flowering phenology, shortening breeding cycles and eliminating field costs associated with maintaining and phenotyping non-selected males. This extends the logic already demonstrated in kiwifruit and date palm, where Y-linked markers eliminated multi-year grow-outs for sex identification (Torres *et al*., 2018; Akagi *et al*., 2019), but goes further: *CsMSFL1* enables within-sex selection for phenological timing, not merely sex determination. In cannabinoid production systems, where pollen release from any retained or undetected male triggers seed set and reduces cannabinoid yield, stratifying males by their *CsMSFL1* genotype also identifies which individuals present the highest risk of early, undetected flowering and thus should be prioritized for removal.

Collectively, this study demonstrates that the genetic architecture of cannabis agronomic traits extends well beyond the autosomes and that male plants, far from being genomic bystanders, carry sex-chromosome-encoded variation of direct quantifiable breeding relevance. The identification of *CsMSFL1* as a near-deterministic regulator of male flowering initiation, *CsWSL1* as an X-linked controller of stem biomass, and *CsMSML1* as a pleiotropic PAR hub coordinating male plant morphology provides the first genomic toolkit for rational paternal selection in dioecious cannabis breeding programs: *CsMSFL1* enables seedling-stage prediction and selection of male flowering phenology; *CsWSL1* offers an X-linked target for stem biomass improvement exploitable in female-mediated recurrent selection; and *CsMSML1* constitutes a co-dominant PAR marker for simultaneous selection of height, stem diameter, branching, and relative growth rate in male plant improvement programs. More broadly, the sex-stratified analytical framework developed here, integrating SNP GWAS, *k*-mer association, and haplotype-resolved sex chromosome assemblies, offers a transferable model for agronomic genomics in any dioecious crop where the genetic contribution of the male sex has been systematically overlooked — including asparagus, kiwifruit, date palm, hop, spinach, and papaya — and establishes that male genomics is not a peripheral concern but a core component of sex-aware crop improvement.

### Supplementary data

**Fig. S1.** Sex chromosome identification based on differential sequencing coverage between male and female accessions.

**Fig. S2.** Sex-biased SNP density along the cannabis X and Y chromosomes.

**Fig. S3.** Sex-differentiation index (FST) along the cannabis sex chromosomes before and after *XYRefiner* filtering.

**Fig. S4.** Pairwise genetic distance heatmaps across cannabis sex chromosome compartments.

**Fig. S5.** Linkage disequilibrium (*r*²) decay across sex-chromosomal compartments in cannabis.

**Fig. S6.** Nucleotide diversity and Tajima’s *D* across cannabis sex chromosomes by compartment and sex stratum.

**Fig. S7.** Combined *k*-mer GWAS Manhattan plots for the whole-genome run.

**Fig. S8.** Combined *k*-mer GWAS Manhattan plots for sex-stratified runs.

**Fig. S9.** *K*-mer GWAS pleiotropic signal hotspots across cannabis sex chromosomes.

**Fig. S10.** Quantile–quantile plots for SNP-based GWAS on cannabis ChrX and ChrY.

**Fig. S11**. Allelic effect plots for genome-wide significant SNP markers on cannabis ChrX.

**Fig. S12.** Allelic effect plots for genome-wide significant SNP markers on cannabis ChrY.

**Fig. S13.** Haplotype block structure for genome-wide significant SNP markers on cannabis ChrX.

**Fig. S14.** Haplotype block structure for genome-wide significant SNP markers on cannabis ChrY.

**Table S1.** Cannabis seed samples from 25 native Iranian populations collected across five Köppen–Geiger climatic zones.

**Table S2.** List of phenological and morphological traits investigated in this study.

**Table S3.** Read-depth ratio summary by chromosomal compartment used for PAR/nonPAR boundary determination.

**Table S4.** Variant counts at each quality control stage across chromosomal compartments in XYRefiner.

**Table S5.** *K*-mer GWAS parameter optimization results for ChrX females across three representative traits.

**Table S6.** Number of significant *k*-mer hits at minimum count thresholds of 1, 5, and 10.

**Table S7.** SNP distribution, LD decay, and haplotype block characteristics across cannabis sex chromosome compartments.

**Table S8.** Number of significant *k*-mer GWAS signal windows per agronomic trait and run.

**Table S9.** Annotated *k*-mer GWAS signal windows on cannabis ChrX and ChrY across all runs.

**Table S10.** Sex-stratified *k*-mer GWAS signal counts per trait and chromosomal region on ChrX and ChrY.

**Table S11.** Linkage disequilibrium among significant *k*-mers per agronomic trait and run on cannabis sex chromosomes.

**Table S12.** Genome-wide significant SNP markers on cannabis sex chromosomes and their putative candidate genes

### Author Contributions

M.B, Conceptualization; methodology; investigation; data curation; formal analysis; resources; visualization; writing – original draft; writing – review & editing. D.T, Conceptualization; supervision; funding acquisition; resources; writing – review & editing.

## Data Availability

Raw genotyping-by-sequencing (GBS) data have been deposited in the NCBI Sequence Read Archive under BioProject accession PRJNA1404300. Processed data supporting the findings are publicly accessible via Figshare (DOI, 10.6084/m9.figshare.32561223). The XYRefiner and XY_kmer_GWAS pipelines are available at github.com/Mehdibabaeii/XYRefiner and github.com/Mehdibabaeii/XY_kmer_GWAS respectively.

## Abbreviations

**BLINK**,: Bayesian-information and linkage-disequilibrium iteratively nested keyway;
*CsMSFL1*: Male-SpecificFloweringLocus1;
*CsMSML1*: Male-SpecificMorphologyLocus1;
*CsWSL1*: StemWeightLocus1;
*DR*^2^: dosage *R*-squared;
F_ST_: fixation index;
GBS: genotyping-by-sequencing;
GWAS: genome-wide association study;
HB: haplotype block;
HWE: Hardy–Weinberg equilibrium;
HXR: hemizygous X region;
LD: linkage disequilibrium;
MAC: minimum allele count;
MAF: minor allele frequency;
MSY: male-specific Y region;
NJ: neighbour-joining;
PAR: pseudoautosomal region;
PVE: phenotypic variance explained.

