## Supplementary Figures S1-S14 for "Sex chromosome-aware GWAS reveals XY-linked agronomic architecture and breeding targets in cannabis"

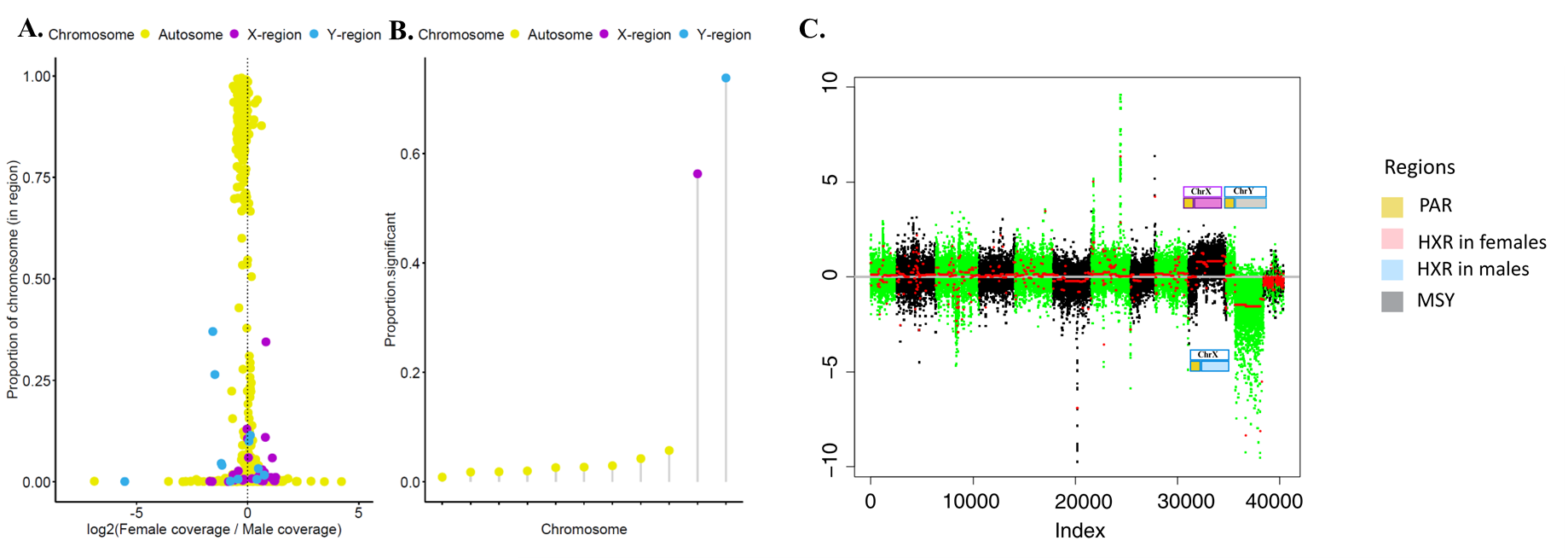

**Figure S1:** Sex chromosome identification in cannabis based on differential sequencing coverage. (**A**) Distribution of log₂(female/male) read-depth ratios per chromosomal region (autosomes, X-chromosomal, Y-chromosomal windows). (**B**) Per-chromosome proportion of windows showing significant female-to-male coverage differences. (**C**) Genome-wide log₂(female/male) coverage ratio across all indexed windows (autosomes: indices 1–31,075; ChrX: 31,076–34,776; ChrY: 34,777–38,427).

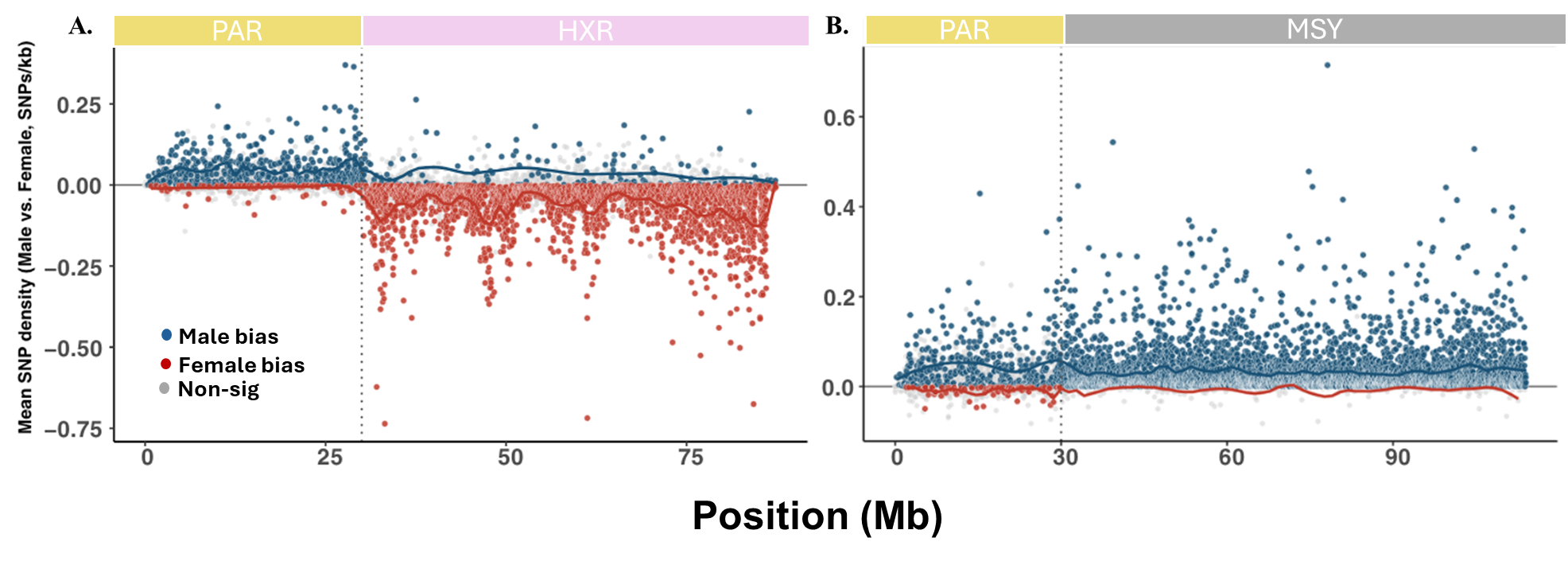

**Figure S2:** Sex-biased SNP density along the cannabis X and Y chromosomes. Mean SNP density difference (Male − Female, SNPs/kb) plotted across genomic positions for (**A**) ChrX and (**B**) ChrY. Points represent individual windows colored by association: male-biased (blue), female-biased (red), or non-significant (gray). Smoothed trend lines highlight regional patterns. Chromosomal domains — pseudoautosomal region (PAR), hemizygous X region (HXR), and male-specific Y region (MSY) — are indicated by colored bars above each panel, with the PAR boundary marked by a vertical dashed line.

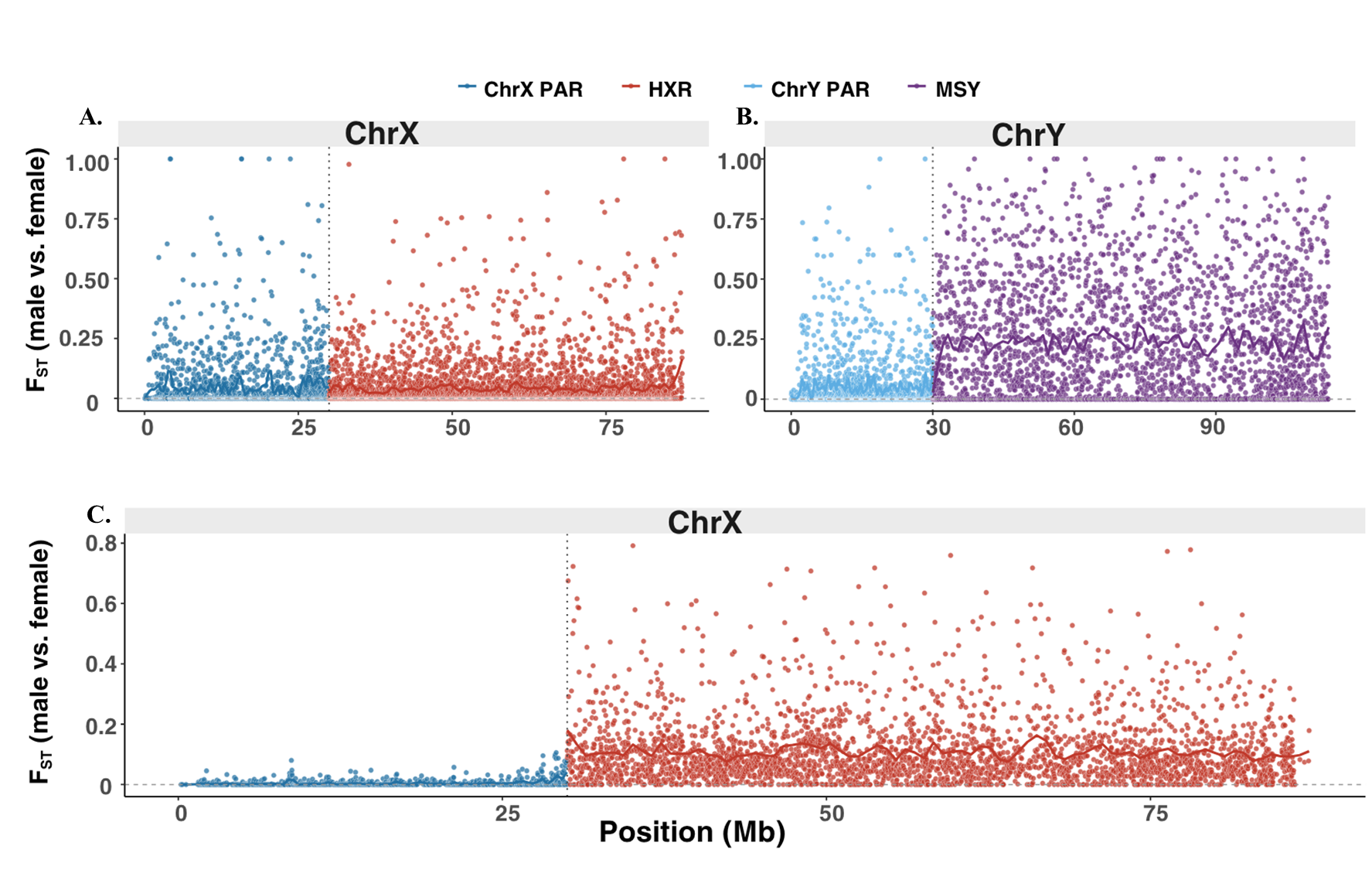

**Figure S3:** Sex-differentiation index (F_ST_) along the cannabis sex chromosomes. Per-window F_ST_ (male vs. female) plotted across genomic positions for (**A**) ChrX, (**B**) ChrY, and (**C**) ChrX shown at a refined scale. Points are colored by chromosomal region: ChrX PAR (dark blue), HXR (red), ChrY PAR (light blue), and MSY (purple). Smoothed trend lines indicate regional averages. The vertical dashed line denotes the PAR boundary. The horizontal dashed line at F_ST_ = 0 serves as a reference baseline.

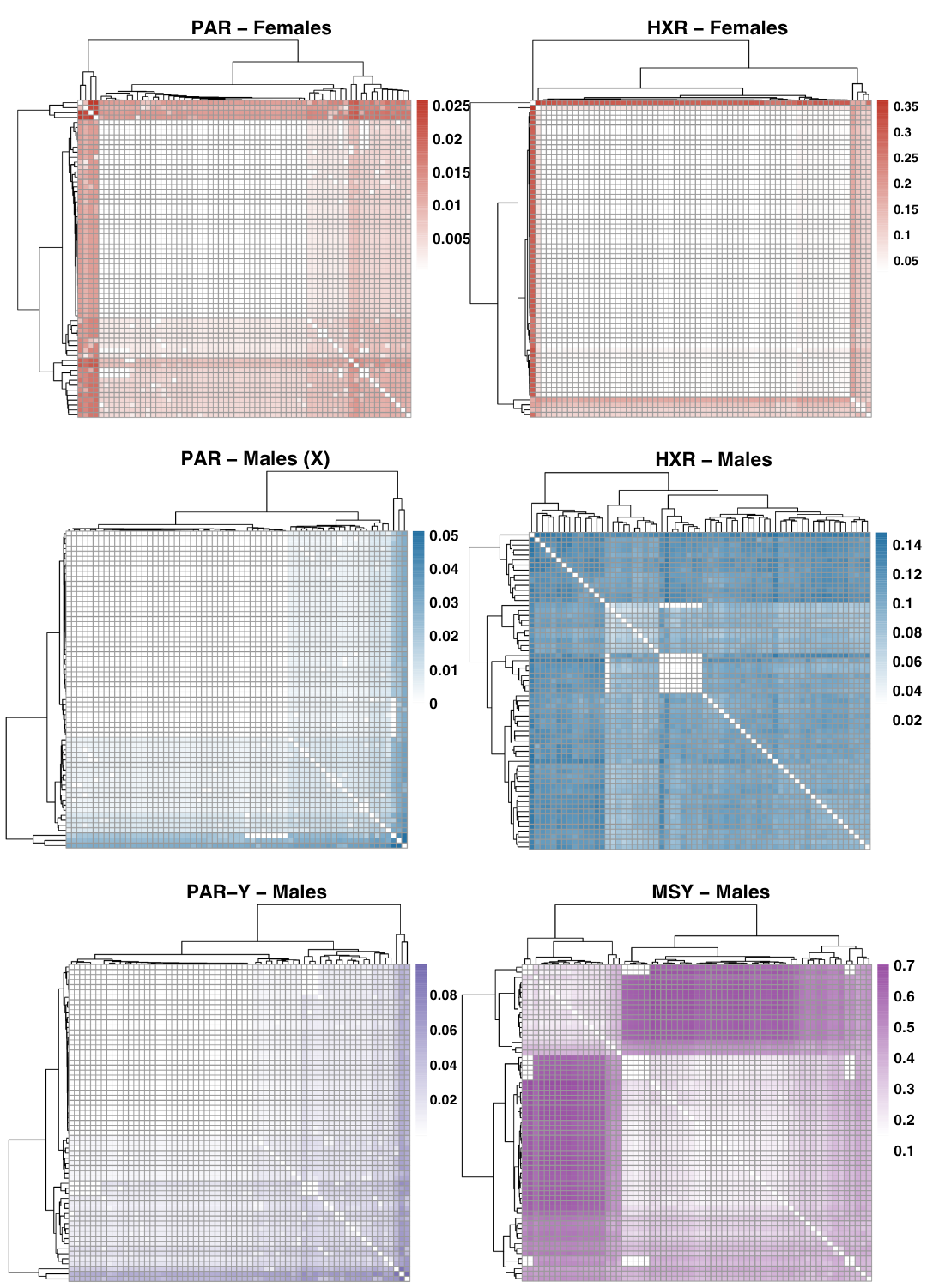

**Figure S4:** Pairwise genetic distance heatmaps (p-distance) between accessions for cannabis sex chromosome compartments: ChrX PAR (females), ChrX PAR (males), ChrX HXR (females), ChrX HXR (males), ChrY PAR (males), and ChrY MSY (males). Samples are annotated by biological sex. Color intensity corresponds to genetic divergence, with darker shades indicating greater distance.

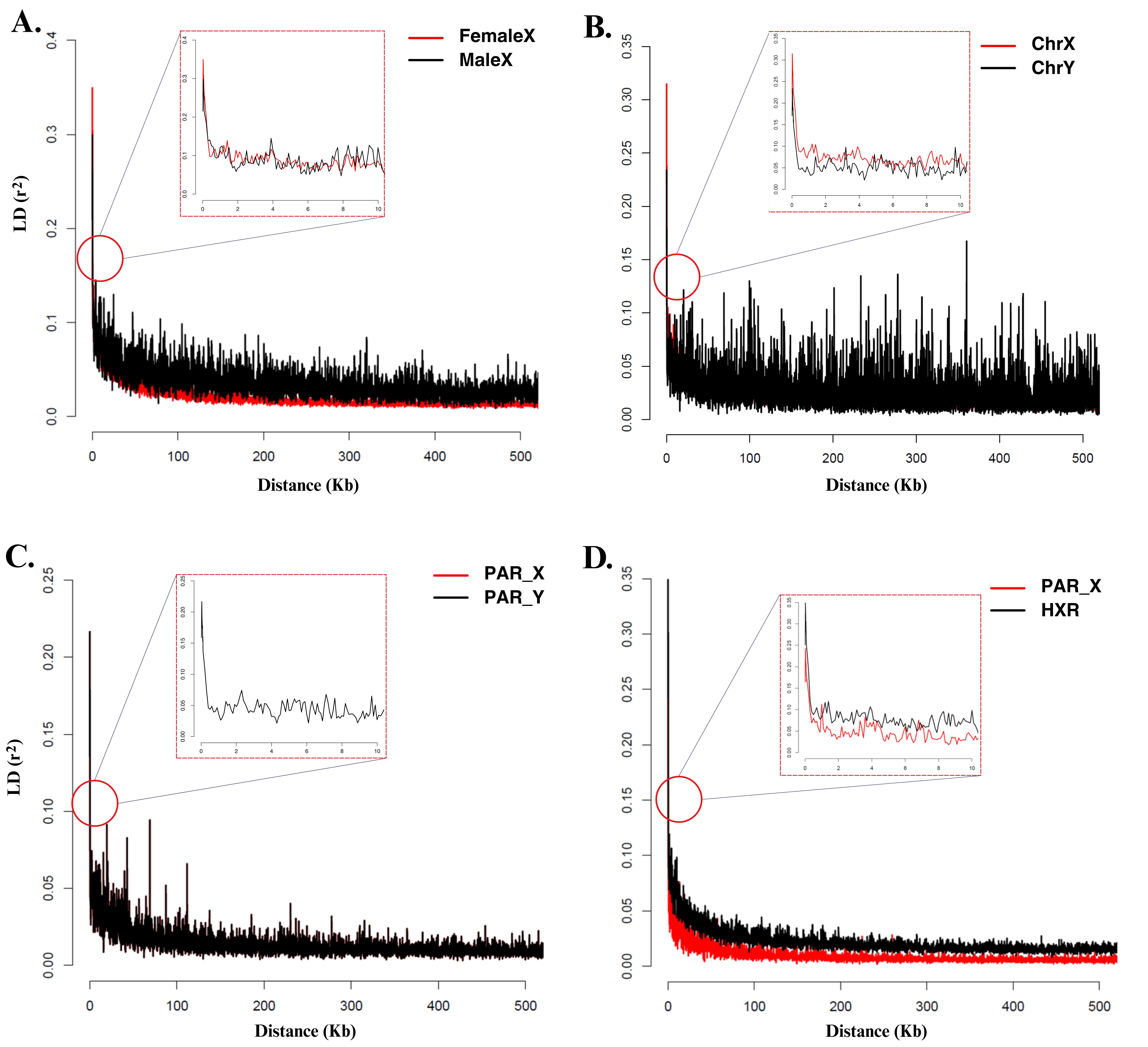

**Figure S5:** Linkage disequilibrium (r²) decay across sex-chromosomal compartments in cannabis. r² plotted as a function of inter-marker distance (kb); insets show expanded 0–10 kb range. (**A**) Female vs. male ChrX. (**B**) ChrX vs. ChrY. (**C**) ChrX PAR vs. ChrY PAR. (**D**) ChrX PAR vs. HXR.

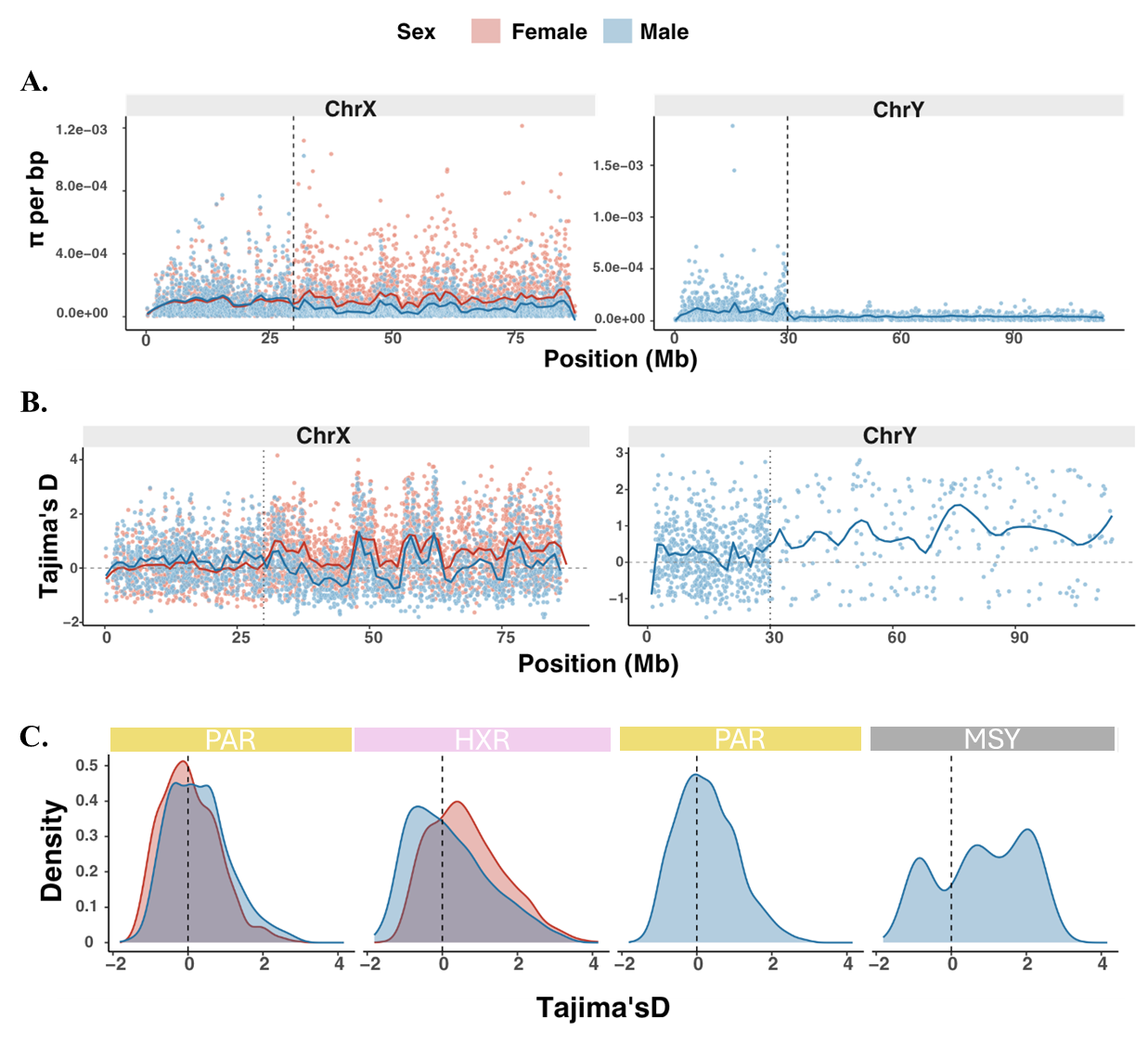

**Figure S6:** Nucleotide diversity and Tajima’s D across cannabis sex chromosomes. (**A**) Windowed nucleotide diversity (θπ; 10 kb windows) plotted along ChrX (left) and ChrY (right) for female (red) and male (blue) samples; vertical dashed line at 30 Mb denotes the PAR/nonPAR boundary. (**B**) Windowed Tajima’s D along ChrX and ChrY by sex stratum; horizontal dashed line at D = 0. (**C**) Density distribution of Tajima’s D values across chromosomal compartments (PAR, HXR, PAR_Y, MSY) for female (red) and male (blue) samples; vertical dashed line at D = 0.

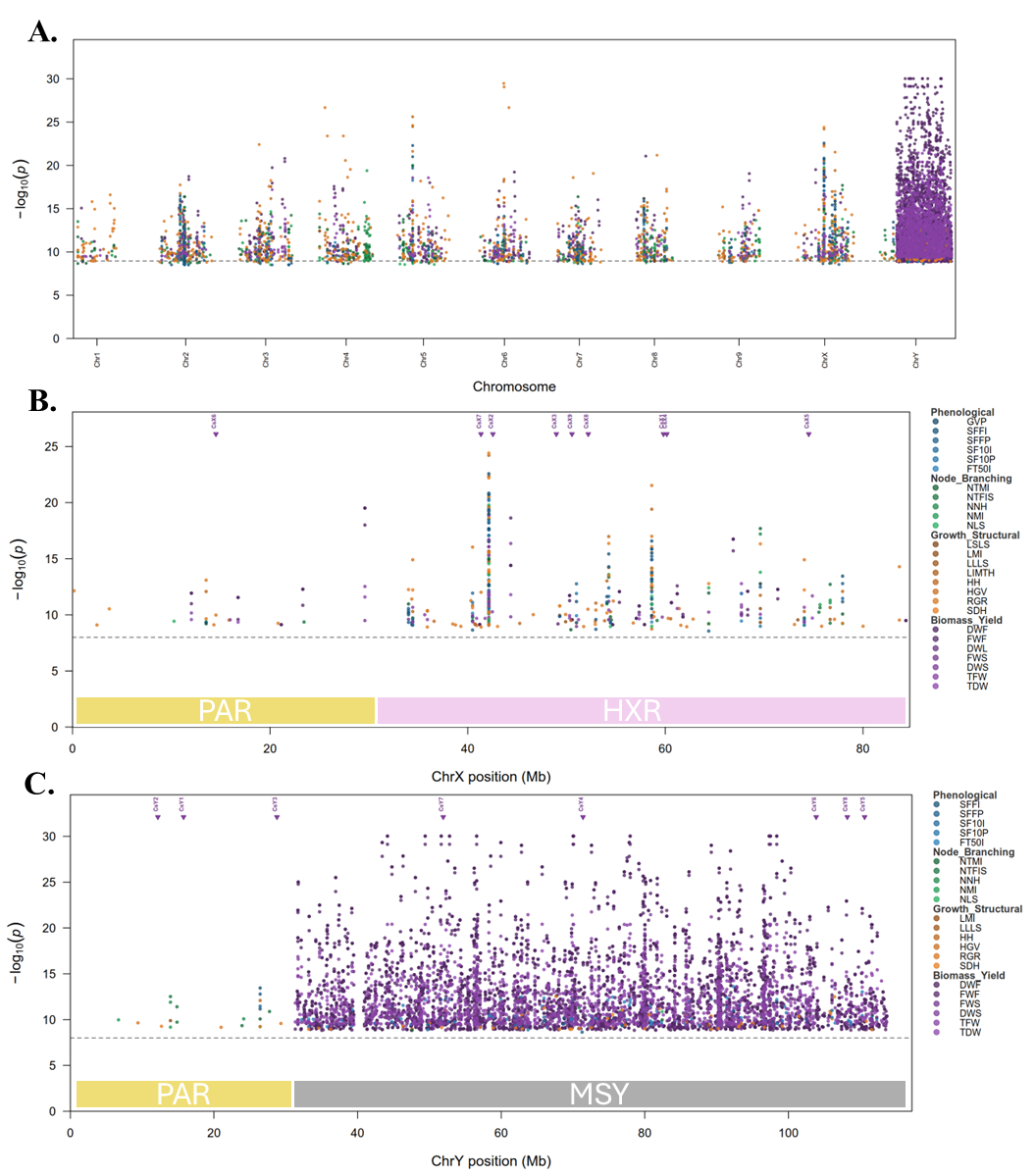

**Figure S7:** Combined k-mer genome-wide association study (GWAS) Manhattan plots for the whole-genome run (n = 145 accessions; 100.3M k-mers). (**A**) Genome-wide view across all 11 autosomes and sex chromosomes; chromosomes colour-coded. (**B**) ChrX signals. (**C**) ChrY signals. Horizontal dashed line: permutation-based significance threshold (α = 0.05). On sex chromosomes, position 30 Mb denotes the PAR/nonPAR boundary (PAR: 0–30 Mb; HXR on ChrX and MSY on ChrY: >30 Mb). Trait abbreviations follow Supplementary Table S2.

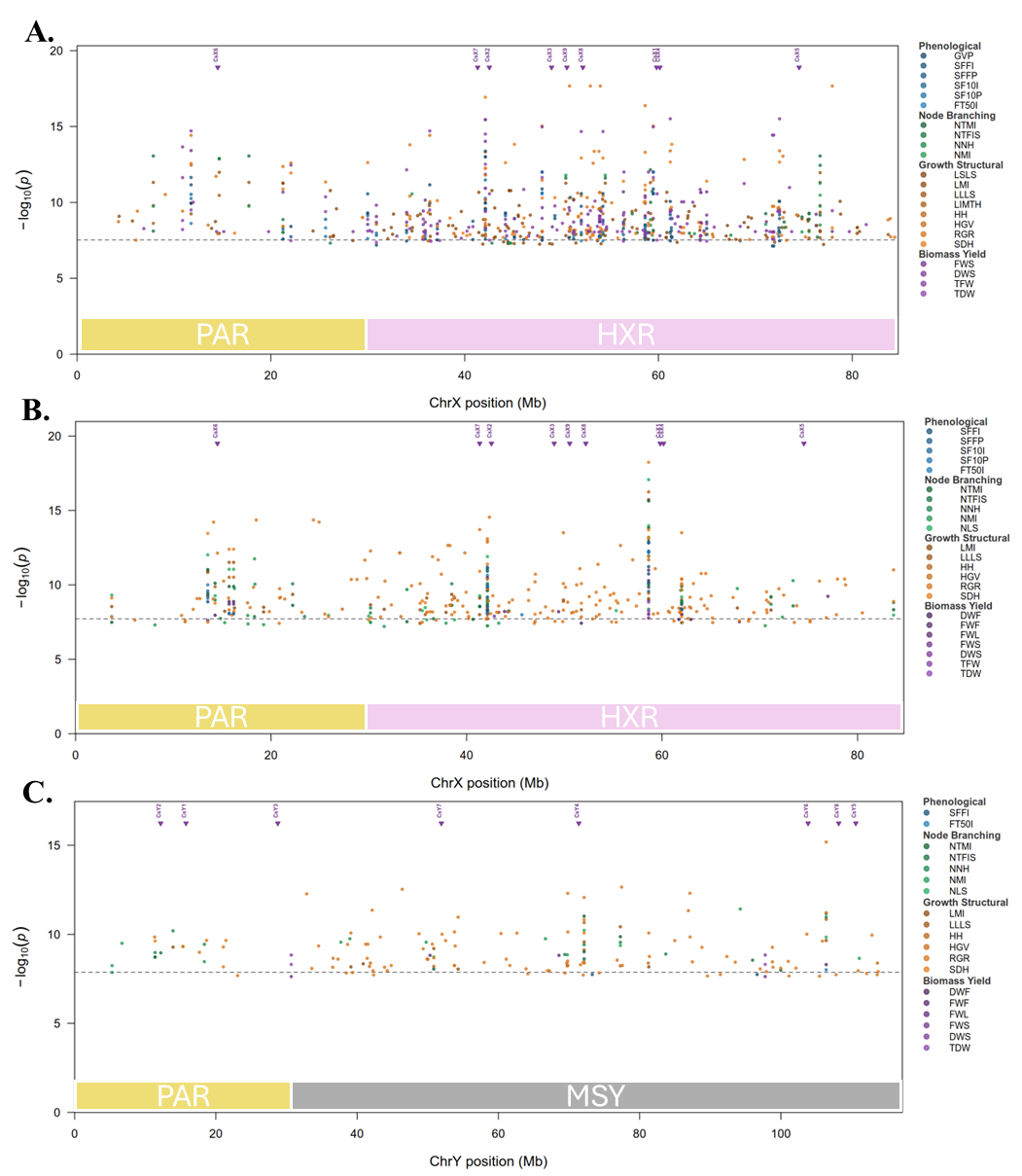

**Figure S8:** Combined k-mer genome-wide association study (GWAS) Manhattan plots for sex-stratified runs. (**A**) ChrX_females (n = 73; 6.74M k-mers). (**B**) ChrX_males (n = 72; 5.87M k-mers). (**C**) ChrY_males (n = 72; 7.41M k-mers). Horizontal dashed line: permutation-based significance threshold (α = 0.05). Position 30 Mb denotes the PAR/nonPAR boundary (PAR: 0–30 Mb; HXR on ChrX and MSY on ChrY: >30 Mb). Trait abbreviations follow Supplementary Table S2.

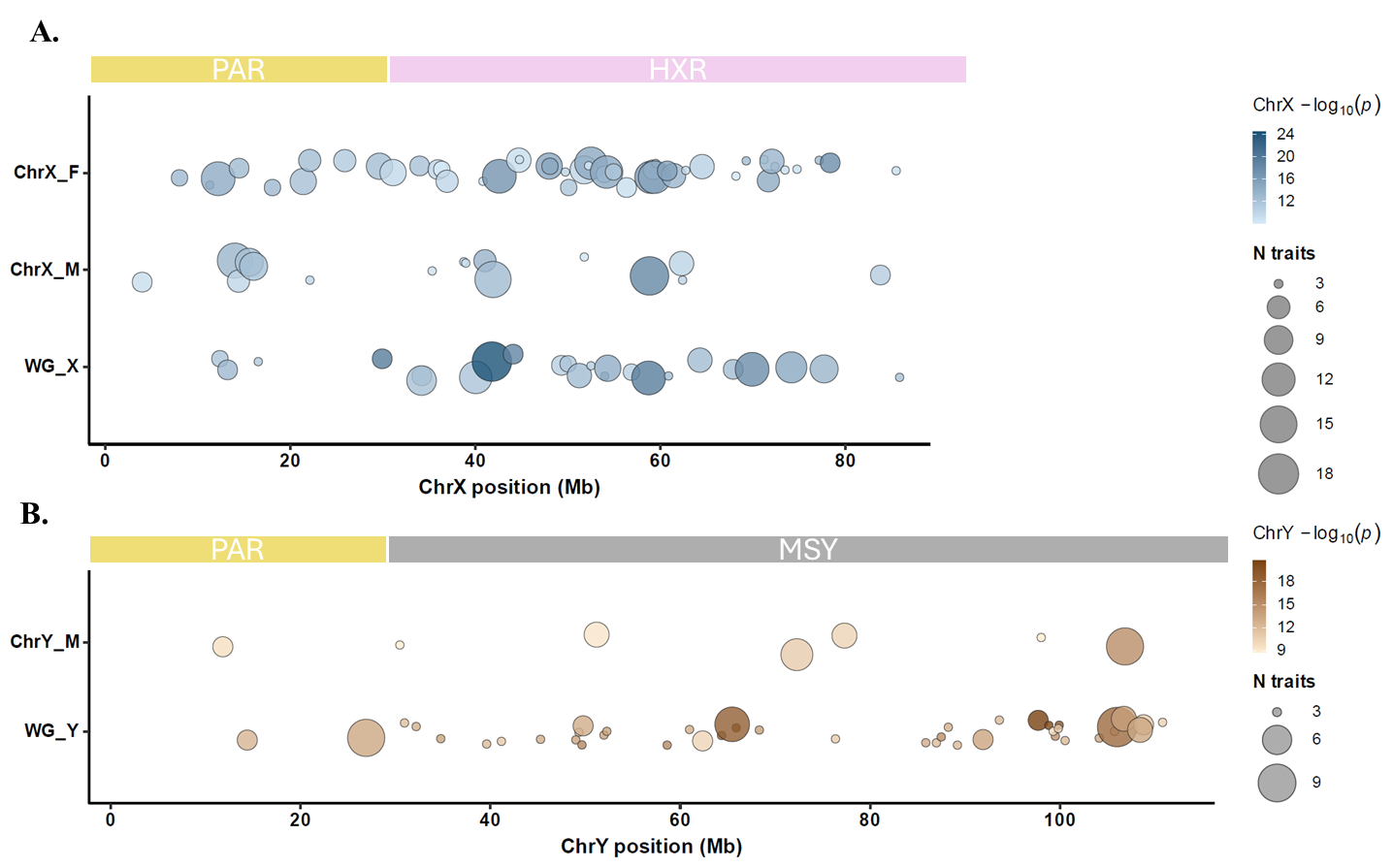

**Figure S9:** K-mer genome-wide association study (GWAS) pleiotropic signal hotspots across cannabis sex chromosomes. Each circle represents a unique genomic window (≤ 250 kb); circle size = number of associated traits (N traits ≥ 3); fill colour = maximum −log_10_ (p) within the window. (**A**) ChrX signals from ChrX_females (ChrX_F), ChrX_males (ChrX_M), and whole-genome ChrX (WG_X) runs. (**B**) ChrY signals from ChrY_males (ChrY_M) and whole-genome ChrY (WG_Y) runs. Position 30 Mb denotes the PAR/nonPAR boundary (PAR: 0–30 Mb; HXR on ChrX and MSY on ChrY: >30 Mb).

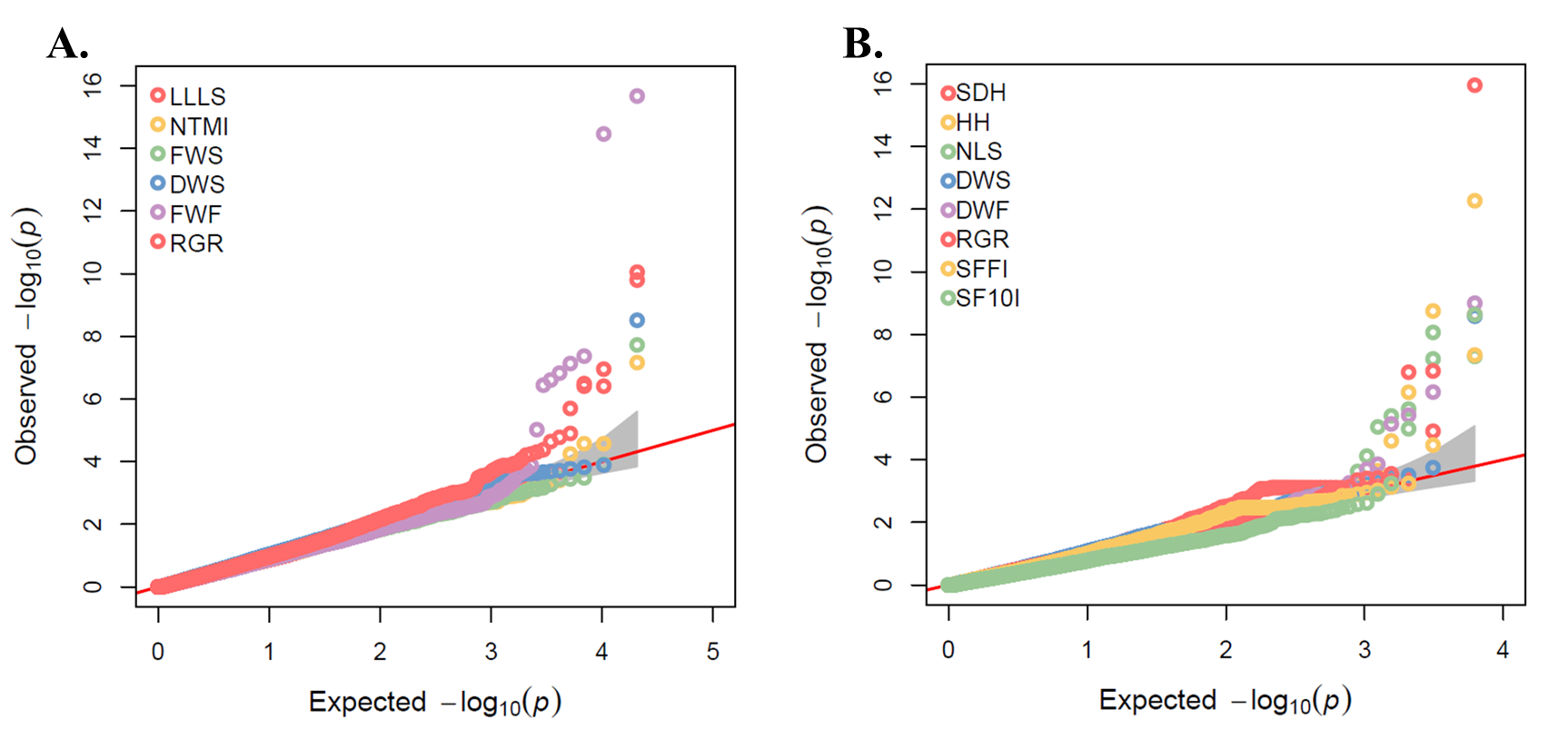

**Figure S10:** Quantile–quantile (QQ) plots for SNP-based genome-wide association study (GWAS) on cannabis sex chromosomes. Observed −log_10_ (*p*) plotted against expected values under the null hypothesis. (**A**) ChrX GWAS (*n* = 145 accessions). (**B**) ChrY GWAS (*n* = 72 males). GWAS performed using BLINK. Trait abbreviations follow Supplementary Table S2.

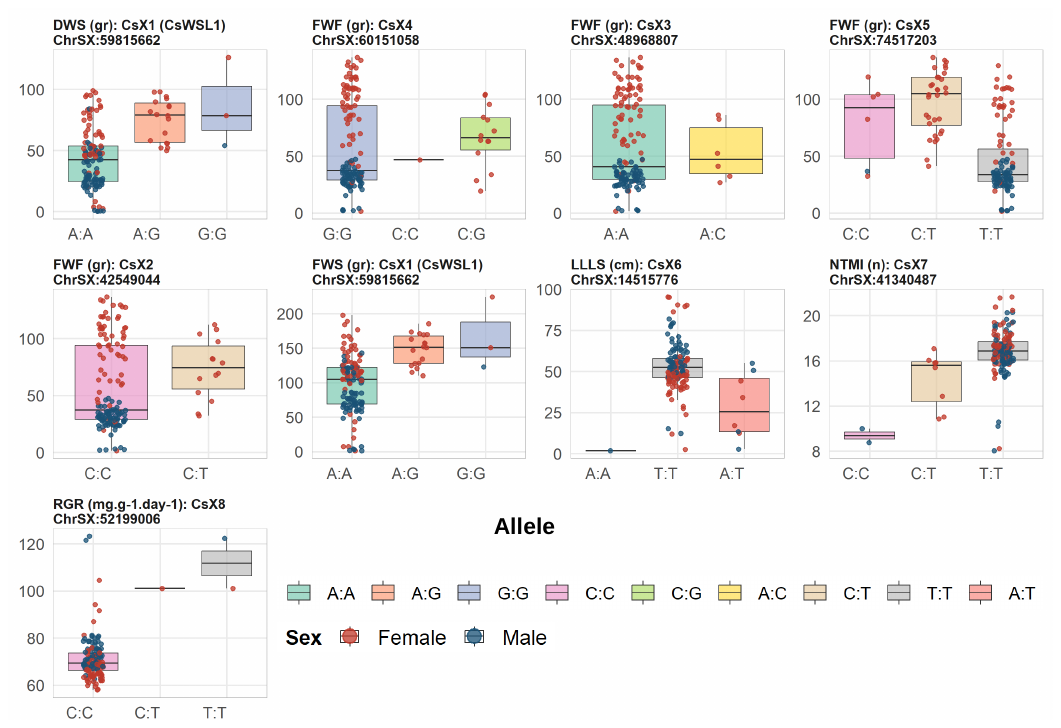

**Figure S11:** Allelic effect plots for genome-wide significant single nucleotide polymorphism (SNP) markers on cannabis ChrX. Each panel shows the distribution of trait values across genotype classes for a specific marker; individual sample points are coloured by biological sex (female: red; male: blue). Markers shown: CsX1 (DWS, FWS), CsX2–CsX5 (FWF), CsX6 (LLLS), CsX7 (NTMI), CsX8 (RGR). Marker details in Table 1 and Supplementary Table S12. Trait abbreviations follow Supplementary Table S2.

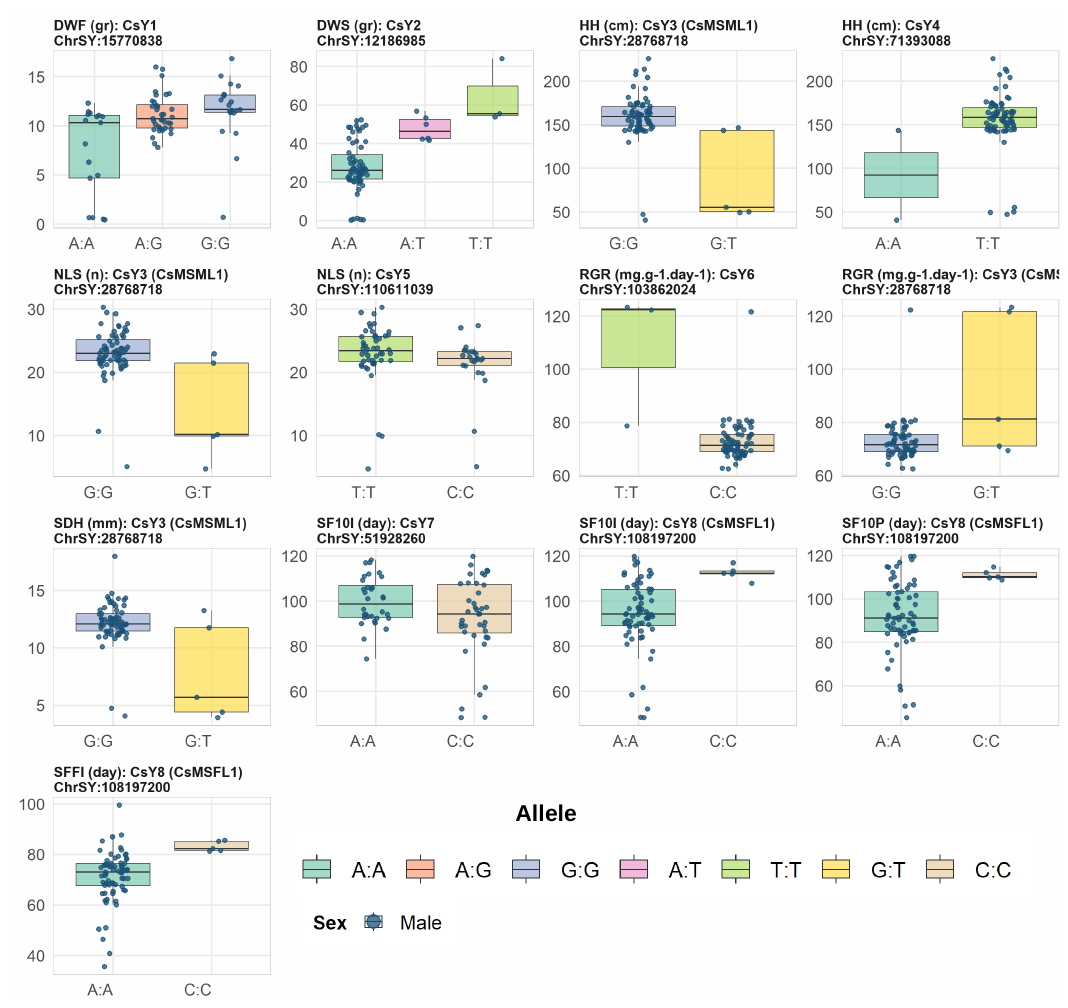

**Figure S12:** Allelic effect plots for genome-wide significant single nucleotide polymorphism (SNP) markers on cannabis ChrY. Each panel shows the distribution of trait values across genotype classes for a specific marker; individual sample points are coloured by biological sex (male: blue). Markers shown: CsY1 (DWF), CsY2 (DWS), CsY3 (HH, NLS, RGR, SDH), CsY4 (HH), CsY5 (NLS), CsY6 (RGR), CsY7 (SF10I), CsY8 (SF10I, SF10P, SFFI). Marker details in Table 1 and Supplementary Table S12. Trait abbreviations follow Supplementary Table S2.

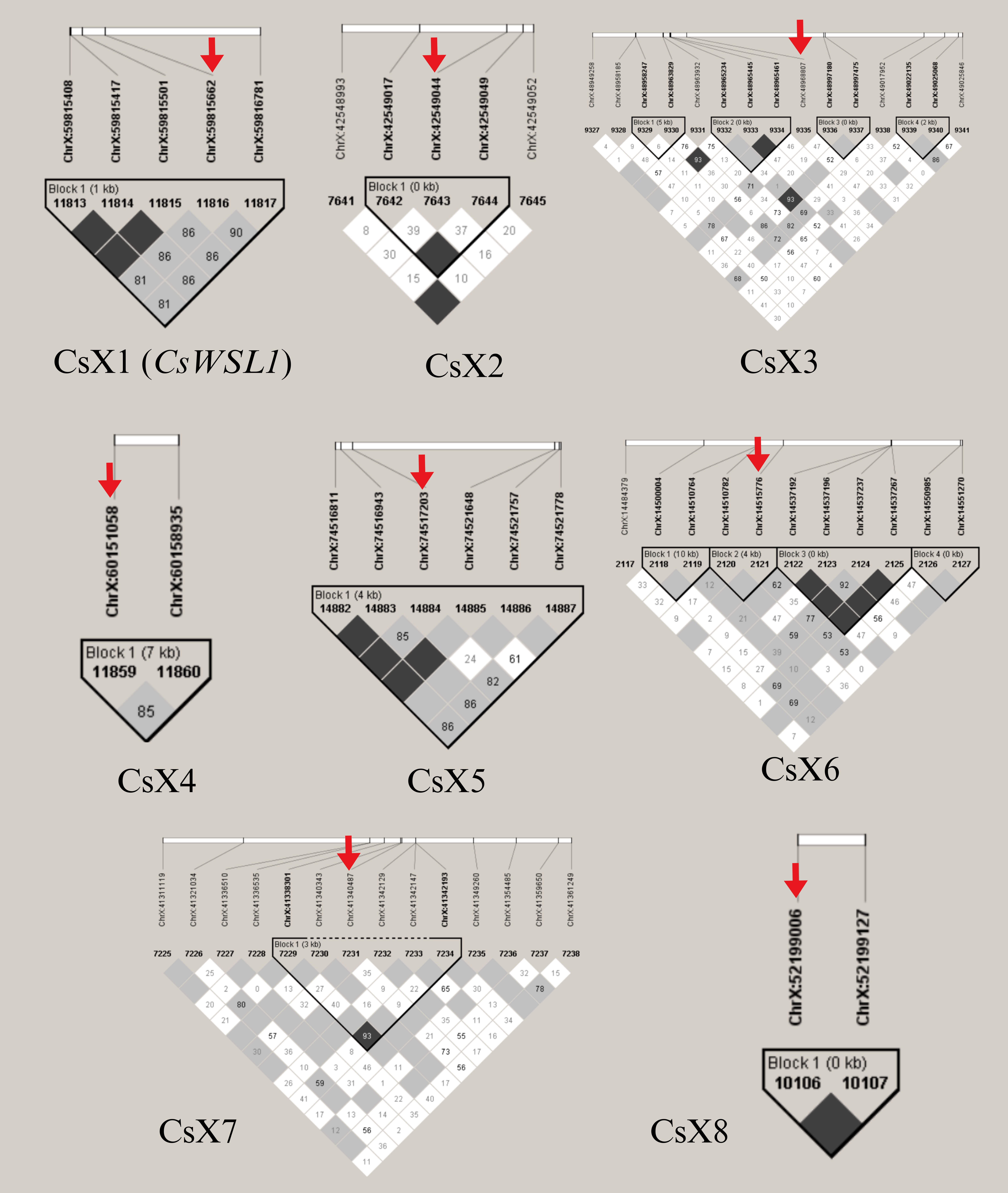

**Figure S13:** Haplotype block structure for genome-wide significant single nucleotide polymorphism (SNP) markers on cannabis ChrX. Pairwise linkage disequilibrium (*r*²) plotted for SNPs flanking each marker; colour scale: white = 0, black = 1; haplotype block boundaries indicated by triangles. Red arrows indicate the position of each significant marker. Marker details in Table 1 and Supplementary Table S12.

**Figure S14:** Haplotype block structure for genome-wide significant single nucleotide polymorphism (SNP) markers on cannabis ChrY. Pairwise linkage disequilibrium (*r*²) plotted for SNPs flanking each marker; colour scale: white = 0, black = 1; haplotype block boundaries indicated by triangles. Red arrows indicate the position of each significant marker. Extended block spans at MSY markers (234–493 kb) consistent with suppressed recombination in the male-specific Y region. Marker details in Table 1 and Supplementary Table S12.
